# Bispecific GD2 x B7-H3 Antibody Improves Tumor Targeting and Reduces Toxicity while Maintaining Efficacy for Neuroblastoma

**DOI:** 10.1101/2024.05.23.595588

**Authors:** Amy K. Erbe, Arika S. Feils, Alina Hampton, Zachary T Rosenkrans, Mildred Felder, Jessica Wiwczar, Daniel J. Gerhardt, Mark Bercher, Belinda Wenke, Callie Haertle, Mackenzie Heck, Sabrina N. VandenHeuvel, Lizzie Frankel, Megan Nielsen, Dan Spiegelman, Noah Tsarovsky, Jen Zaborek, Alexander L. Rakhmilevich, Jacquelyn A. Hank, Eduardo Aluicio-Sarduy, Jonathan W. Engle, Jonathan H. Davis, Bryan Glaser, Vladimir Subbotin, Roland Green, Reinier Hernandez, Bonnie Hammer, Paul M. Sondel

## Abstract

The current treatment for neuroblastoma involves an immunotherapy regimen that includes a monoclonal antibody that recognizes disialoganglioside (GD2), expressed at high levels on neuroblastoma. GD2 is not present on most normal tissues but is expressed on nerves. Thus, anti-GD2 treatment causes substantial, dose-limiting, neuropathic pain. B7-H3 is overexpressed on multiple tumor types, including neuroblastoma, with minimal normal cell expression and is absent on nerves. We designed a bispecific antibody (bsAb) that requires simultaneous binding of these two tumor antigens to achieve tight-binding of tumor cells. Our preclinical research shows that when compared to an anti-GD2 monospecific antibody, the GD2xB7-H3 bsAb has improved tumor specificity with similar efficacy and reduced toxicity. Since this bsAb does not bind to nerves, it may be possible to administer increased or additional doses beyond the tolerable dose of monospecific anti-GD2 antibodies, which could improve therapeutic efficacy and quality of life for patients with neuroblastoma.

---

Neuroblastoma is an extracranial solid tumor of the sympathetic nervous system that affects ∼800 US children annually. Nearly half of newly diagnosed patients present with high-risk neuroblastoma, for which the standard multimodal treatment plan is divided into 3 phases: 1) Induction (multi-agent chemotherapy and surgery) to reduce the measurable disease; 2) Consolidation (myeloablative chemotherapy, autologous stem cell transplant, radiation therapy) targeting primary and persistent disease sites; 3) Maintenance (immunotherapy and differentiating agent therapy) to eradicate any residual disease.^1^ The current standard of care immunotherapy regimen given during the maintenance phase includes dinutuximab, a chimeric monoclonal antibody (mAb) that recognizes the GD2 disialoganglioside, along with granulocyte-macrophage colony-stimulating factor (GM-CSF).^1, 2, 3^ Preclinical data and clinical genotyping studies suggest dinutuximab acts largely through antibody-dependent cell-mediated cytotoxicity (ADCC) involving NK cells, neutrophils and monocytes/macrophages.^4, 5, 6, 7^ In a randomized Phase III clinical trial, dinutuximab significantly improved overall and progression-free survival rates.^3, 8^ Regimens using different or variant anti-GD2 mAbs (e.g., naxitimab, dinutuximab-beta or hu14.18K322A) also show comparable antitumor benefit.^9, 10, 11^

While minimally expressed on most normal tissues, GD2 is expressed on nerves, resulting in severe neuropathic pain felt by nearly all patients receiving anti-GD2 therapy.^3, 12, 13^ This transient neuropathic pain was the major determinant of the maximum tolerated dose (MTD) for dinutuximab in Phase 1 studies, causing the current dinutuximab dose regimen to be 17.5mg/m^2^/day, corresponding to ∼5-10% of the dose of other tumor-reactive FDA approved mAbs that also act via ADCC, such as rituximab or cetuximab.^14, 15^ Furthermore, three recent clinical studies of anti-GD2 mAb for neuroblastoma showed that higher serum levels of anti-GD2 mAb were associated with better antitumor outcome.^10, 16, 17^ This suggests that administration of greater dosing of anti-GD2 mAbs (similar to that of other therapeutic antibodies), if tolerated, may improve antitumor outcome.

To address this, we developed a bispecific human IgG antibody that has one antigen-binding fragment (Fab) arm specific for GD2 and the other Fab arm specific for B7-H3 (CD276). B7-H3, also overexpressed on neuroblastomas, has minimal expression on normal cells, and is not expressed on nerves.^18, 19^ The fucosylated form of this GD2xB7-H3 bispecific Ab (GD2xB7-H3 bsAb) is designated INV721, and its afucosylated form is INV724 (see below). The monomeric affinity of each arm is only weak to moderate, leading to inefficient binding to cells expressing only one of the two targets. Tight cell surface binding of INV721 (or INV724) is driven by avidity, requiring simultaneous recognition of both GD2 and B7-H3. Thus, it should bind well to neuroblastomas that express both targets, while not binding to nerves that only express GD2 but not B7-H3. By enhancing the tumor-specificity with this GD2xB7-H3 bsAb, and eliminating nerve binding, we should substantially reduce the painful side-effects that neuroblastoma patients experience during their anti-GD2 therapy.

## Results

### INV721 *In Vitro* Antigen/Target Specificity

Attempts have been made to reduce the toxicities associated with anti-GD2 mAb therapy and/or augment its activity, including incorporating species variations (human, mouse, or chimeric mAbs) and altering Fc regions (to reduce complement activation and reduce fucosylation).^2, 20, 21, 22, 23, 24^ While some of these modifications resulted in a partial reduction in clinical toxicities, including neuropathic pain, patients still experienced pain during treatment.^2, 13^ INV721 was uniquely designed using Invenra’s B-Body^®^ technology^255^ to optimize the neuroblastoma-targeting specificity, with individual Fab components that recognize the target antigens monovalently, such that strong avidity is achieved only on cells co-expressing both GD2 and B7-H3 (**Fig 1**). To create the optimal bsAb, several separate Fab binders of varying affinities for B7-H3 and GD2 were tested combinatorially in a bispecific format and assessed for binding to cells expressing GD2 and/or B7-H3 (data not shown). The toxic side effcts associated with the use of dinutuximab are largely linked to the on target/off tumor GD2-targeting to nerve cells, which are bound much more readily by dinutuimxab (monovalent Kd to GD2=10nM)^26^ vs. the bsAb (monovalent Kd to GD2>1µM). The GD2xB7-H3 bsAb retains strong binding to tumor cells which co-express both GD2 and B7-H3, as the B7-H3 arm has moderate affinity (Kd=2nM) and thus the bsAb has stronger avid binding when both the GD2 and B7-H3 arms engage (**Fig. 1**).

**Figure 1:**
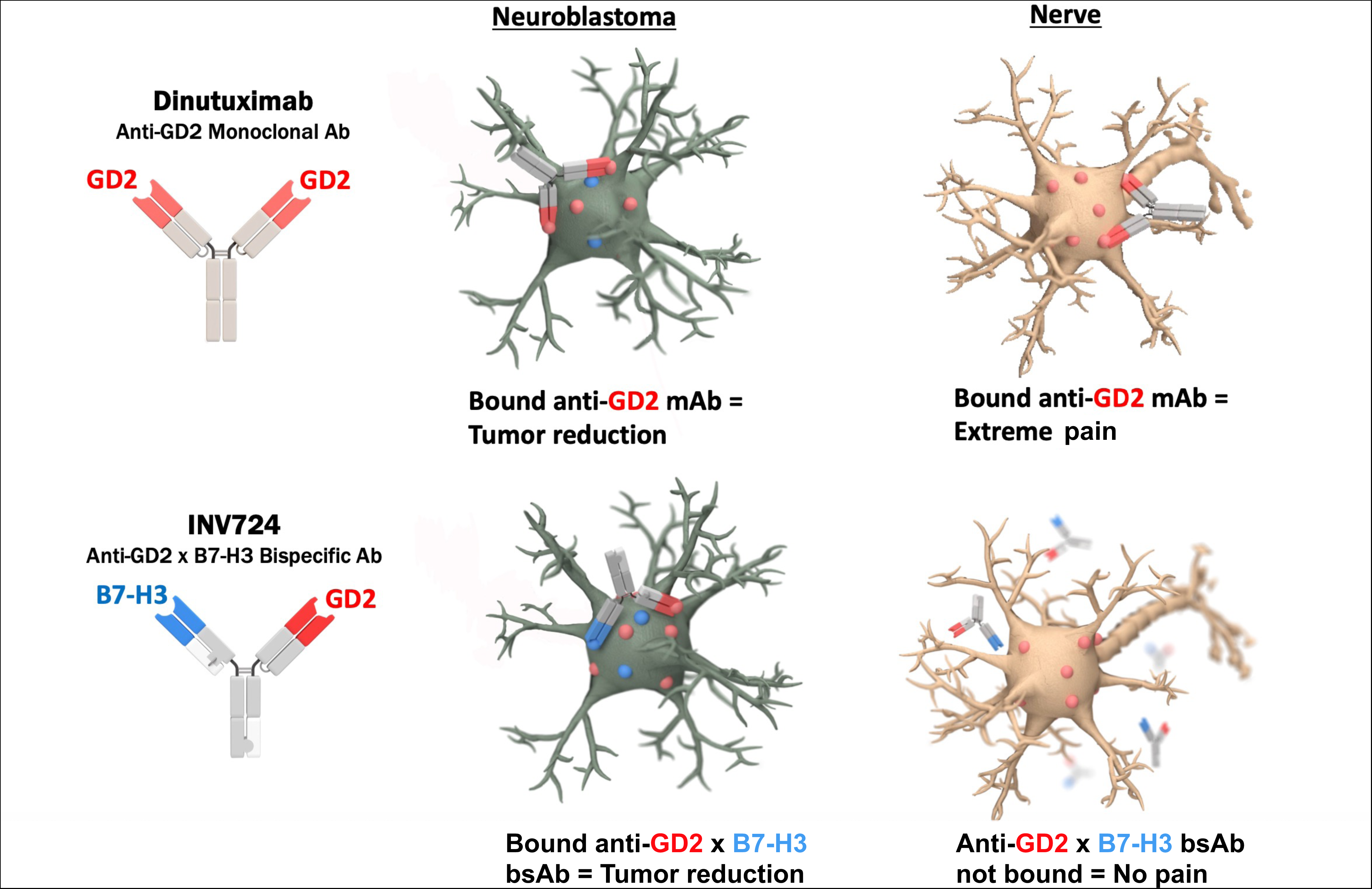
INV721 has improved tumor specificity and has high avidity only when both Fab arms are bound to target antigens. Dinutuximab, with each Fab arm recognizing GD2, can bind to both GD2+ tumors as well as GD2+ nerves (top panel). INV721, with one Fab arm specific to GD2 and the other to B7-H3, can only bind with high-avidity to neuroblastoma tumors (expressing both GD2 and B7-H3) and cannot bind to nerves, which lack B7-H3 expression (bottom panel).

To separate the affinity of the individual Fab arms of INV721 from the avidity of the combination, we created single Fab-arm versions of INV721: 1) the B7-H3 Fab with a non-binding Fab arm (B7-H3xNon-Binding) or 2) the GD2 Fab with a non-binding Fab arm (GD2xNon-Binding). Using B78 murine melanoma cells that express GD2 with or without co-expression of human B7-H3 (i.e., GD2^+^/B7-H3^+^ simulating tumor, or GD2^+^/B7-H3^-^ simulating nerve), we tested the cell-binding affinity of these one-armed variants compared to the INV721 bsAb. When both target antigens are co-expressed on target tumor cells (**Fig. 2A**), INV721 binds strongly, driven by avidity, even at low concentrations, but the B7-H3xNon-Binding Ab shows binding only at higher concentrations (displaying moderate-affinity monovalent binding of the anti-B7-H3-Fab). GD2xNon-Binding Ab binding was not detected, even at high concentrations, demonstrating the inability of the monovalent anti-GD2 arm to bind well on its own (**Fig 2A**). In contrast, on cells expressing only GD2 but not B7-H3, as seen on nerves (**Fig. 2B**), none of these 3 antibodies bind, even at high concentrations, reflecting the need for INV721 to recognize both GD2 and B7-H3 on the same cell. Against the human neuroblastoma cell line, CHLA20, which endogenously expresses both GD2 and B7-H3, we observe increased avidity with INV721 as compared to the B7-H3xNon-Binding Ab, and we observe no binding of the GD2xNon-Binding Ab (**Fig 2C**).

**Figure 2.**
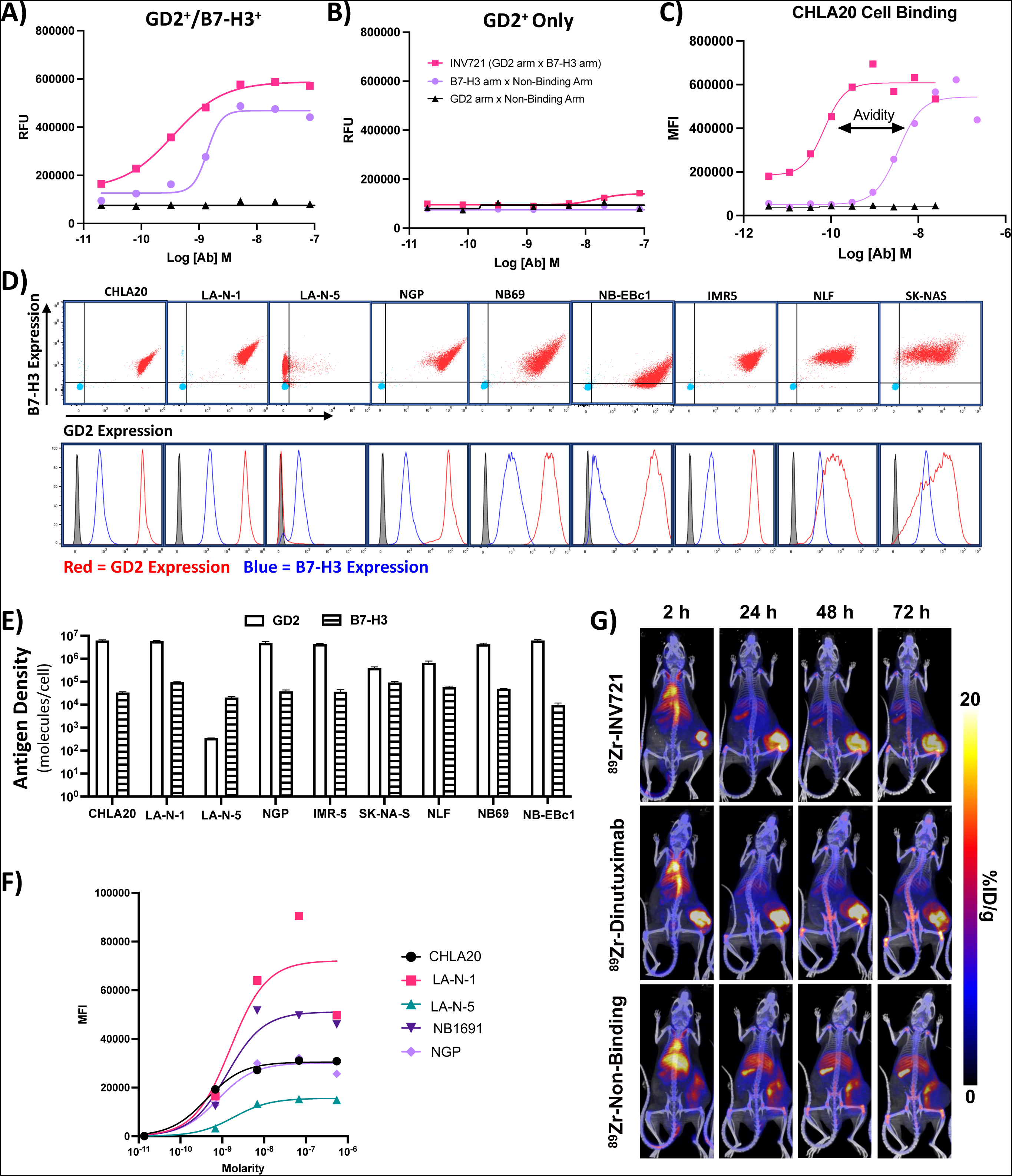
INV721 binds neuroblastoma target cell. INV721 with B7-H3 Fab x GD2 Fab arms, or antibody variations with the GD2 Fab arm x a non-binding arm or the B7-H3 Fab arm x a non-binding arm were tested by flow cytometry for binding to B78 melanoma cell line variants (GD2^+^/B7-H3^+^) in **A** or (GD2^+^/B7-H3^-^) in **B**. **A)** Stronger binding is observed to the GD2^+^/B7-H3^+^ cells when both B7-H3 Fab x GD2 Fab arms can bind (INV721, pink line) vs. when only one Fab arm can bind (B7-H3 Fab x Non-Binding Fab arm, purple line or GD2 Fab x Non-Binding Fab arm, black line). **B)** Very little binding is observed for the INV721 B7-H3 Fab x GD2 Fab arms in cells that only express the GD2 target (GD2^+^/B7-H3^-^). **C)** Human CHLA20 neuroblastoma cells which express endogenous GD2 and B7-H3 bind with higher avidity to INV721 with B7-H3 Fab x GD2 Fab arms (pink) than to B7-H3 Fab arm x a non-binding arm (purple), and no binding was observed to CHLA20 from the GD2 Fab x Non-Binding Fab arm (black). **D)** Nine human neuroblastoma cell lines tested here demonstrate their level of expression for B7-H3 and GD2 as shown by flow cytometry. The 2-dimensional dot-plots are shown in the top set of graphs (with B7-H3 on Y axis and GD2 on X-axis), and the histograms are shown in the bottom panel, with GD2 in red and B7-H3 in blue, and the secondary antibody alone shown in gray. **E)** For 8/9 human neuroblastoma cell lines (all but LA-N-5), the GD2 antigen density is greater than the antigen density of B7-H3, but all 9 lines express detectable levels of both antigens. **F)** INV721 titration testing shows 5/5 human neuroblastoma cell lines tested bind INV721. **G)** NRG mice bearing an NGP tumor in the right flank show ^89^Zr-INV721 is taken up and bound to the tumor 2 hrs after injection and is retained within the tumor for 72 hrs, similar to that of ^89^Zr-Dinutuximab. No tumor uptake was observed for the ^89^Zr-Non-Binding Ab (which does show non-specific uptake in liver and spleen).

As noted previously, Majzner et al. showed that GD2 and B7-H3 are co-expressed on most human neuroblastoma lines (**Fig 2D**), with GD2 usually expressed at a higher antigen density than B7-H3 (**Fig. 2E**).^27^ All human neuroblastoma cell lines tested bound to INV721, with variations in the observed amount of binding that may be dependent on the antigen density ratio of GD2 to B7-H3 (**Fig. 2F**). Notably, the LA-N-5 cell line expressed B7-H3 with low but detectable GD2 (**Fig. 2D-E**) and was still capable of binding to INV721 (**Fig 2F**).

To test *in vivo* tumor targeting, we assessed *in vivo* trafficking of INV721 using positron emission tomography (PET) imaging of Zirconium-89 (^89^Zr) radiolabeled mAbs injected intravenously (IV) in NRG mice bearing human NGP neuroblastoma xenografts (which endogenously express B7-H3 and GD2). We found that ^89^Zr-INV721 targets, binds to and is retained in established NGP tumors at levels comparable to ^89^Zr-dinutumab (**Fig 2G**).

### *In Vitro* Immune-Mediated Antitumor Efficacy

A major mechanism of action of anti-GD2 mAbs is to elicit ADCC against GD2^+^ cells.^28, 29, 30^ ADCC involves cell-mediated killing by Fc gamma receptor (FC_Y_R)-bearing immune cells that recognize the Fc portions of mAbs bound to the tumor cell surfaces. To augment ADCC, it may be beneficial to utilize mAbs that are not internalized, or are internalized slowly, after binding to the tumor cell surface, thus maintaining external exposure of the Fc portion of the mAb. Prior studies have reported that anti-GD2 mAbs, including dinutuximab, internalize within 16-24hrs once bound to target cells, which may serve as a mechanism of immune resistance.^31^ We also observed internalization of dinutuximab over 16hrs when tested on five separate human tumor cell lines that endogenously express GD2 and B7-H3. In contrast, we found minimal internalization of INV721 as well as anti-B7-H3 mAb, I7-01, in this same assay (**Fig. 3A**). Despite its internalization, dinutuximab is still capable of eliciting strong ADCC efficacy at 50ng/ml (**Fig. 3B**), with similar efficacy as that of INV721, which has the same human IgG1 backbone as dinutuximab (**Fig 3B**).

**Figure 3:**
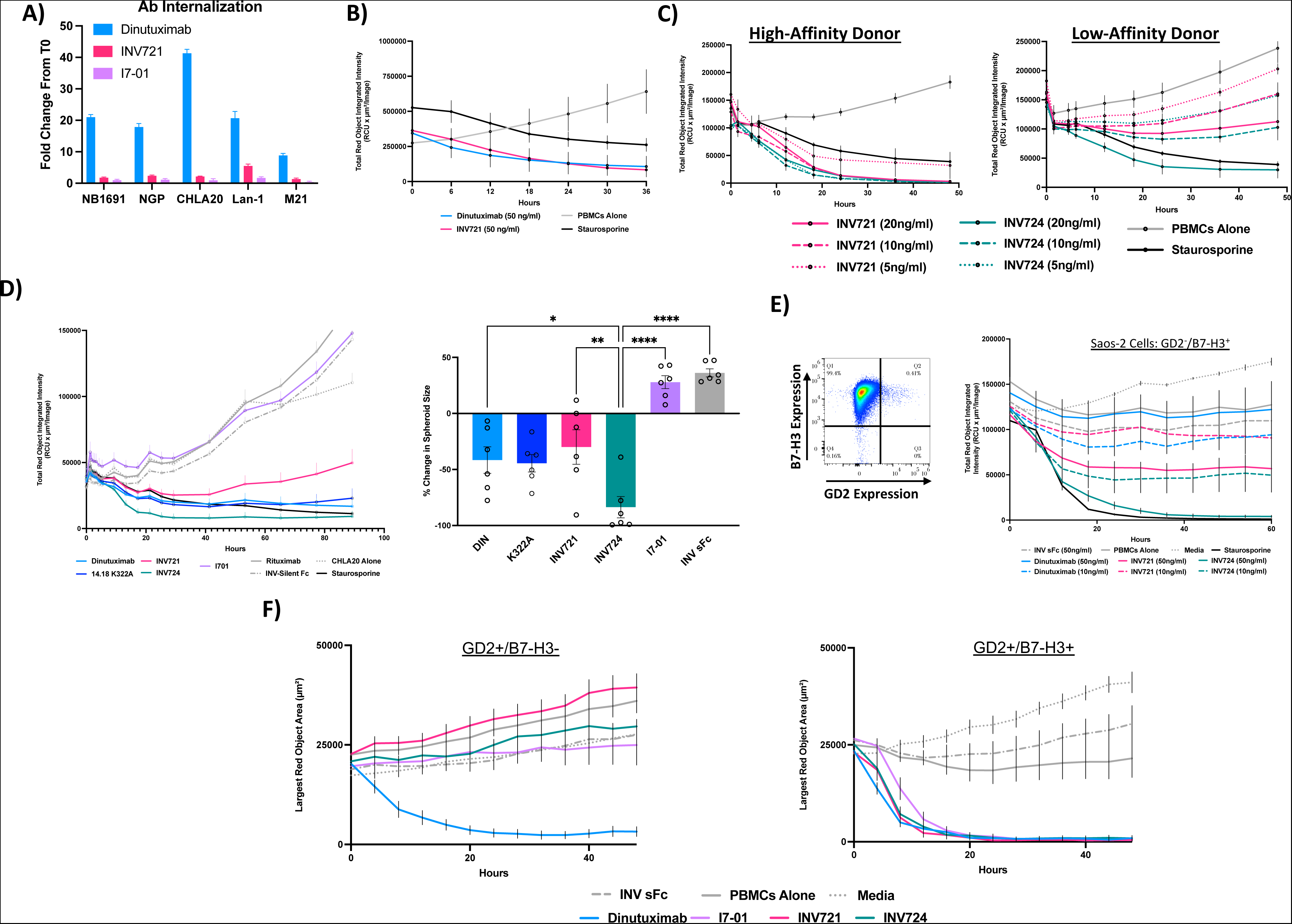
GD2^+^/B7-H3^+^ bispecific antibody efficiently induces ADCC. A) For five different human cell lines tested (neuroblastoma cell lines NB1691, NGP, CHLA20, and LA-N-1 shown in Fig. 2D, and the melanoma line M21), dinutuximab internalized to a higher degree than INV721 antibody or anti-B7-H3 mAb, I7-01. Results for all 5 cell lines are shown as the ratio of the amount of internalized antibody after 24 hours, divided by the background level of antibody internalized at time-0 (T0), and expressed as “fold change”. **B)** Using an IncuCyte to assess ADCC, human CHLA20 neuroblastoma cells plated as spheroids showed increased spheroid size over time when plated with PBMCs without mAb (gray line); and decrease in spheroid size when plated with staurosporine as a positive cytotoxic agent control (black line). Spheroid size decreased at the same rate when plated with either dinutuximab (blue line) or INV721 (pink line) each at 50ng/ml. This indicates that INV721 is potent at ADCC as dinutuximab. **C)** PBMCs from a donor with a high-affinity FC_Y_Rgenotype exhibit strong ADCC against human CHLA20 spheroids (in an IncuCyte assay) with INV721 that has a standard fucosylated Fc IgG1 end, at concentrations of 10 and 20ng/ml; with decreased killing at 5ng/ml; in contrast killing at all 3 concentrations (5, 10 and 20ng/ml) is potent and comparable using INV724. PBMCs from a donor with a low-affinity FC_Y_R genotype are capable of strong ADCC, but only at the higher concentration tested (20ng/ml) and only with INV724 (i.e., with the afucosylated Fc IgG1 end). **D)** ADCC IncuCyte assays against CHLA20 spheroids were repeated with PBMCs from 6 different healthy donors together with mAbs (each at 40 ng/ml). The left plot shows killing over time; each separate line represents the combined data from the PBMCs from all 6 donors, for the antibody or conditions indicated by the color legend. The right plot shows % change in spheroid size calculated for each donor (as individual circles) as well as the mean % change for all 6 PBMC donors, from the time 0 (when PBMCs, spheroids and Abs were first plated together) to the 24 hrs timepoint for the 6 antibodies shown. Treatment with PBMCs incubated together with CHLA20 with rituximab (gray line, a control, anti-CD20 mAb that does not bind these tumors) resulted in the tumors growing, as did treatment with anti-B7-H3 mAb (I7-01) and treatment with a B7-H3 x GD2 bispecific antibody that has an effector silent Fc “sFc” (INV sFc). The sFc antibody has the same Fab binding regions as INV721 and INV724, but it has mutations in the IgG backbone such that the Fc gamma receptors on immune cells fail to recognize it and cannot mediate ADCC. The sFc antibody serves as a control to show that reduction in spheroid size via INV721 targeting is ADCC-mediated. INV724 treatment resulted in more spheroid shrinkage compared to K322A (monospecific anti-GD2 with afucosylated Fc region), but the difference was not significant. Treatment with dinutuximab and GD2xB7-H3-targeted antibodies (INV721 or INV724) resulted in spheroid shrinkage in nearly all 6 donors; INV724 showed significantly more ADCC capabilities than dinutuximab and INV721 (fucosylated form of the GD2 x B7-H3 bispecific Ab). ****p<0.0001; **p<0.01; *p<0.05. **E)** The human osteosarcoma cell line, Saos-2, expresses B7-H3 but does not express GD2 (left panel). Using PBMCs from a donor with a high-affinity FC_Y_R genotype (right panel), at 50 ng/ml, potent ADCC was achieved with INV724 (green solid line); at the lower INV724 concentration of 10 ng/ml (green dashed line), ADCC was still seen, but decreased. Substantially less ADCC was seen with INV721 or dinutuximab at either concentration. **F)** B78 murine GD2+ melanoma cells expressing both GD2 and human B7-H3 (right plot) are capable of being killed via ADCC when plated together with PBMCs from a donor with high affinity FC_Y_R genotype and INV724 (green line) or dinutuximab (blue line) at 40 ng/ml. Cells expressing only GD2 without human B7-H3 (left plot, simulating nerves) were not killed by ADCC via PBMCs (high-affinity donor) + INV724 (green line), but were killed by PBMCs + dinutuximab (blue line).

We and others have shown that response to mAb immunotherapy can vary based on an individual’s FC_Y_R genotype.^32, 33, 34^ In addition, the ability of FC_Y_Rs to bind to tumor-bound Ab is significantly influenced by the fucosylation patterns of the Ab, whereby reduced fucosylation of the Fc portion of the Ab enhances FC_Y_R binding by 10-100 fold.^35^ We developed the GD2xB7-H3 bsAb with two different Fcs. INV721 has a standard Fc region with an IgG1 backbone that is fucosylated (like dinutuximab). In contrast, INV724 has an afucosylated Fc, which enables augmented binding to FC_Y_Rs on NK cells, enhancing its potential to eradicate tumors through ADCC. Using a high-affinity FC_Y_R donor, we found that INV721 and INV724 have similar ADCC efficacy at higher concentrations (20 and 10ng/ml), while at lower concentrations (5ng/mL) we observe reduced ADCC by INV721 compared to INV724 (**Fig 3C**, left; **Supplemental Fig. 1**). However, for a low-affinity FC_Y_R donor, while INV724 was still capable of eliciting ADCC at 20ng/ml, INV721 had lost its potency (**Fig 3C**, right). Furthermore, using INV724, we observe significantly enhanced antitumor activity against CHLA20 tumor cells via ADCC (vs. dinutuximab or INV721) when comparing the ADCC capabilities from 6 different donors (**Fig. 3D)**. Thus, INV724 became our lead candidate for the *in vivo* efficacy studies. Notably, we see that INV724 also mediates greater ADCC than an anti-B7-H3 mAb created with the anti-B7-H3 binding domain of INV724 (**Fig. 3D**, I7-01).

Even though the higher avidity of INV724 enables potent ADCC killing of GD2^+^/B7-H3^+^ cells but not GD2^+^/B7-H3^-^ cells, the higher affinity of the anti-B7-H3 Fab of INV724 compared to the anti-GD2 Fab (**Fig 2A)** enables the B7-H3 arm of INV724 to bind to GD2^-^/B7-H3^+^ cells with monovalent affinity at higher concentrations. With this binding affinity, INV724 can elicit potent ADCC against B7-H3^+^ tumor cells that do not co-express GD2 (Saos-2) if tested at a concentration of 50ng/ml, but much less so at 10ng/ml (**Fig 3E**). Importantly, we confirmed that this is not the case for the anti-GD2 Fab arm of INV724, which does not allow INV724 to bind to GD2^+^/B7-H3^-^ cells. B78 cells that express only GD2, without co-expression of B7-H3 (simulating the GD2^+^/B7-H3^-^ phenotype of nerve cells) were killed via ADCC by dinutuximab, but not by INV724 (**Fig. 3F**, left graph). In contrast, B78 murine melanoma cells that expressed both GD2 and B7-H3 were killed by both dinutuximab and INV724 via ADCC (**Fig. 3F**, right). Thus, even though both B78 variants express GD2, INV724 cannot kill those without B7-H3 co-expression, even at high concentrations with a high-affinity FC_Y_R donor.

### INV721 *In Vivo* Tumor-Targeting and Specificity

Using our murine B78 melanoma tumor model, we created four expression variants of B78 that do or do not express GD2 or human B7-H3 (i.e., B78-GD2^+^/B7-H3^+^, B78-GD2^-^/B7-H3^+^, B78-GD2^+^/B7-H3^-^ and B78-GD2^-^/B7-H3^-^). We confirmed the antigen density and expression of GD2 and B7-H3 on each variant (**Fig. 4A** and **4B**). We found that INV721 binds with high avidity to the B78-GD2^+^/B7-H3^+^ variant, and with moderate affinity to B78-GD2^-^/B7-H3^+^ cells if at high concentrations of Ab (**Fig. 4C**). Importantly, even at high mAb concentrations, INV721 did not bind to GD2^+^/B7-H3^-^, which simulates the expression of these markers on nerve cells (**Fig. 4C**).

**Figure 4:**
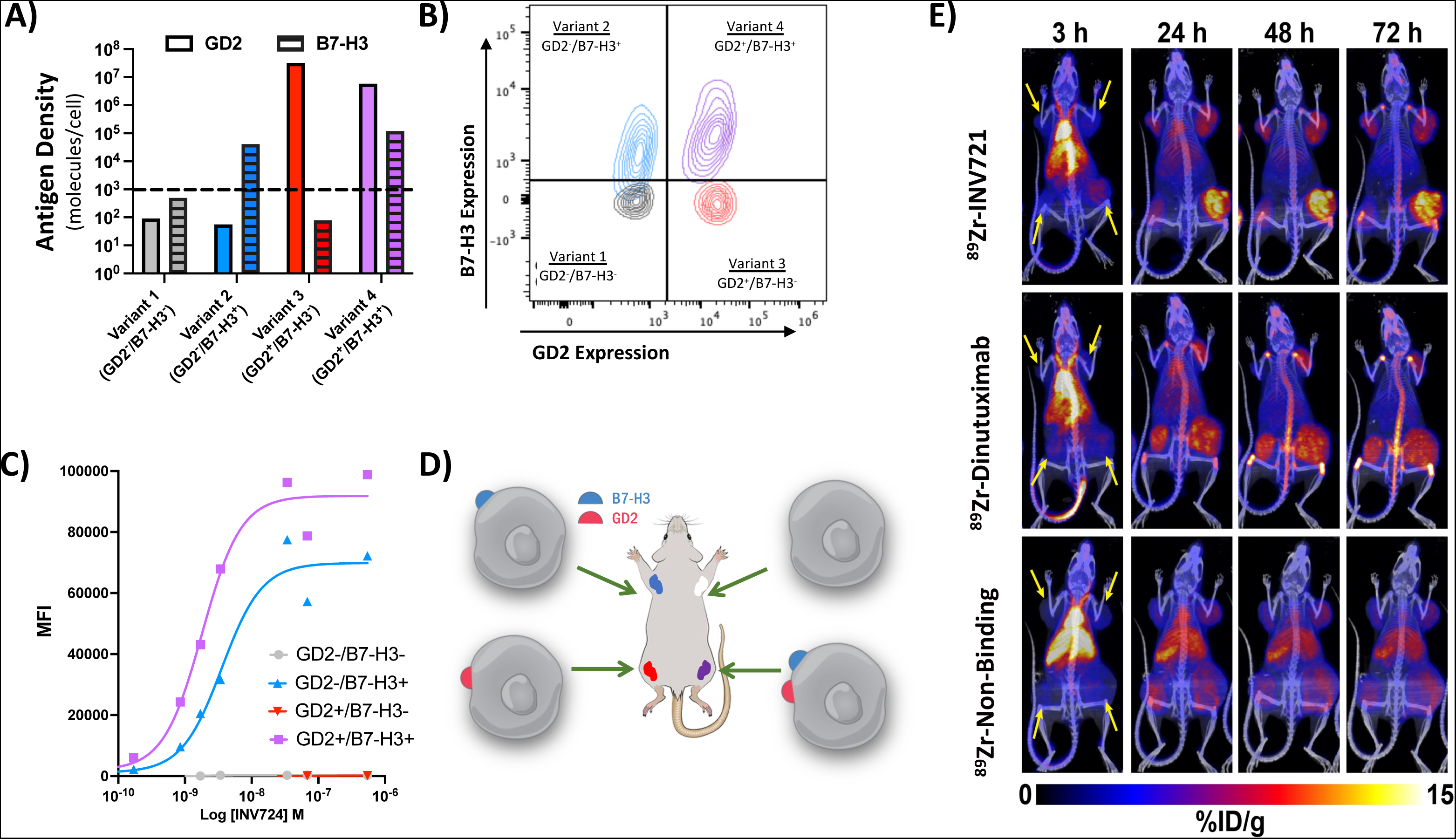
INV721 specifically targets GD2^+^/B7-H3^+^ tumors with high avidity *in vivo.* A) The antigen densities of GD2 and B7-H3, measured as mean number of surface receptors per cell, were determined by flow cytometry for the 4 B78 melanoma cell line GD2/B7-H3 variants with expression variations of GD2 and B7-H3. For each of the 4 variants, the bar on the left (solid color) shows the density of GD2, and the bar on the right (with laddered black lines) shows the density of B7-H3. The dashed horizontal line at a value of ∼10^3^ surface receptors per cell indicates the level of background detection we considered as our cut-off for detectible antigen expression (refer to **Supplemental** Fig. 2). **B)** Contour plots of both B78-GD2 and B7-H3 antigen expression are shown for Variant 1=GD2^-^/B7-H3^-^ (gray), Variant 2= GD2^-^/B7-H3^+^ (blue), Variant 3= GD2^+^/B7-H3^-^ (pink), Variant 4= GD2^+^/B7-H3^+^ (purple). **C)** INV721 binding to these B78 variants depends on the expression of both B7-H3 and GD2. Variant 4 (GD2+/ B7-H3^+^, purple line) showed the highest binding of INV721; Variant 2 (GD2-/B7-H3^+^, blue line) also bound INV721, but only at higher concentrations, indicating less avidity for INV721 than Variant 4. INV721 showed minimal binding to Variants 1 (GD2-/B7-H3^-^, black line) and 3 (GD2+/B7-H3^-^, blue line), which are both superimposed on the X-axis, both of which do not express B7-H3. **D)** Diagram of placement locations within the mice for each of the 4 B78 variants; note the tumor at upper right expresses neither antigen. **E)** Mice with 4 B78 variant tumors (yellow arrows indicate tumors at 3 hr time point) used in PET imaging showing the strongest amount of tumor-uptake of INV721 (top row) when both GD2 and B7-H3 are expressed (bottom right tumor), while minimal uptake is seen in the 3 other tumors: when B7-H3 alone is highly expressed (upper left tumor), when GD2 alone is expressed (bottom left tumor), and when neither GD2 nor B7-H3 are expressed (upper right tumor). Dinutuximab (middle row) showed similar high uptake when GD2 alone is expressed (bottom left tumor) or when GD2 is co-expressed with B7-H3 (bottom right tumor). Minimal uptake of the non-binding control antibody (bottom row) was observed in any of the tumors.

To further confirm the ability of INV721 to traffic specifically to GD2^+^/B7-H3^+^ targets *in vivo*, we developed a “4-tumor model” with each individual mouse injected with the four separate B78 variants each in a different location (**Fig. 4D**), allowing us to monitor the specificity of Ab trafficking based on the differential antigens expressed on each of the four tumors. PET imaging showed that IV-injected ^89^Zr-INV721 targets and binds to GD2^+^/B7-H3^+^ tumors (lower right flank), but not to GD2^+^/B7-H3^-^ tumors (simulating nerve tissue, lower left flank) (**Fig. 4E**). In contrast, as expected, and shown in **Fig 4E**, ^89^Zr-dinutuximab binds strongly to GD2^+^/B7-H3^-^ tumors (simulating nerve tissue, lower left flank) and GD2^+^/B7-H3^+^ tumors (representing clinical neuroblastoma, lower right flank).

### Antitumor *In Vivo* Immunotherapeutic Efficacy

Using syngeneic murine models for both neuroblastoma and melanoma, we observed strong *in vivo* antitumor efficacy of INV724 in mice bearing GD2^+^/B7-H3^+^ tumors. Murine tumor models (B78 melanoma, syngeneic to C57Bl/6 mice and NXS2 neuroblastoma, syngeneic to A/J mice) that express GD2 were transduced to express human B7-H3, and treated with an *in situ* vaccine regimen developed in our lab that includes radiation therapy (RT) together with an antitumor mAb and the stimulatory cytokine, IL-2.^36, 37^ Treatment of mice bearing B78-GD2^+^/B7-H3^+^ tumors (**Fig. 5A-C)** or mice bearing NXS2-GD2^+^/B7-H3^+^ tumors (**Fig 5D-F**) with INV724 (added to RT + IL-2) improved response compared to RT + IL-2 without antibody). Additionally, we demonstrated single agent efficacy with INV724 treatment in nude mice bearing experimental metastases of human CHLA20 neuroblastoma (injected IV), which led to significantly increased progression free survival (PFS), overall survival (OS) and tumor free mice as compared to untreated mice (**Fig 5G-J**). No statistically significant differences were observed between the efficacy of INV724 and dinutuximab in any of these studies (**Fig 5A-J**). These *in vivo* tumor-binding (**Fig. 4E**) and antitumor efficacy studies (**Fig. 5**) demonstrate the potential *in vivo* utility of INV724 to serve as an alternative to dinutuximab in cancer immunotherapy.

**Figure 5.**
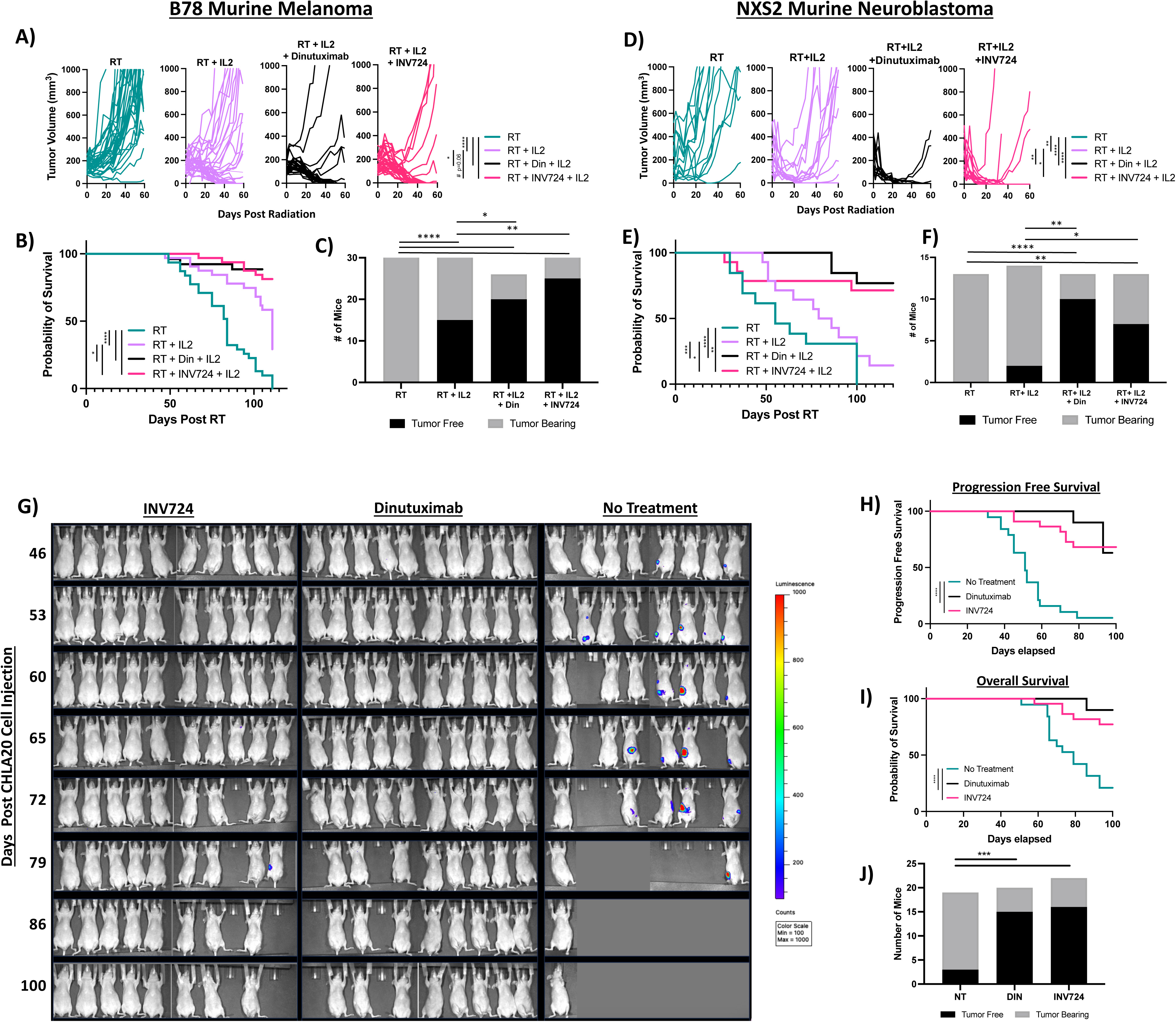
Treatment with INV724 antibody has antitumor efficacy against murine tumors. Mice bearing murine B78 melanoma tumors (**A-C**) or NXS2 neuroblastoma tumors (**D-F**) were randomized to receive external beam radiation [RT (12Gy) D0] +/- IL-2 (75,000 U/dose) +/- Ab (40 µg/dose; either INV724 or Dinutuximab depending on group) D5-D9. **A-C)** C57Bl/6 mice bearing syngeneic B78-B7-H3^+^/GD2^+^ melanoma tumors (n=10-16 mice/group) and **D-F)** A/J mice bearing syngeneic NXS2 neuroblastoma tumors (n=13-14 mice/group) showed that mice treated with RT + INV724 + IL2 (pink) or RT + Dinutuximab + IL2 (black) had reduced tumor growth rate measured after 60 days of growth (**A, D**), significantly improved survival after 125 days (**B, E**) and a significantly increased fraction of mice remaining tumor free (**C, F**) as compared to mice treated with RT + IL2 (purple) or RT alone (green) [as measured on Day 60 (**B**) and Day 90 (**E**)]. Note that mouse deaths due to tumor burden or from tumor-unrelated deaths are included in the overall survival curves. Nude mice were injected via tail vein with human neuroblastoma, CHLA20-luciferase cells on D0, then treated either with INV724 (25 µg/dose, n=22) or dinutuximab (25 µg/dose, n=20) or PBS control (n=19) via retro-orbital IV injection on D1, D4, D7, D10 (**G-J**). **G)** Mice were monitored weekly via an *in vivo* imaging system (IVIS) for tumor development, and all images were normalized to the same color scale (range 100 to 1000) for uniformity. **H**) PFS was calculated based on IVIS signal and survival: if no visible signal was detected by IVIS or distended abdomen, and if mice remained alive. **I**) Survival was also determined for each group. **J)** A significant increased fraction of mice remaining tumor free (as of Day 100) was observed in mice treated with INV724 or dinutuximab compared to mice not treated. Figures A-C, D-F, and G-J each present the combined results of 2 similar, separately performed experiments, and indicate statistical comparisons between groups showing significance. ****p<0.0001; ***p<0.001; **p<0.01; *p<0.05.

### *Ex Vivo* Nerve Binding and *In Vivo* Pain Toxicity Studies

Besides tumor expression, GD2 is also expressed on human central nervous system cells and the myelin sheaths of peripheral nerves.^38, 39^ GD2 expression on peripheral nerves is implicated in the peripheral neuropathic pain experienced by patients during dinutuximab therapy.^40^ To confirm that Fab arms of this GD2xB7-H3 bsAbs (INV721 and INV724 have the same Fab regions) are incapable of binding to nerves, we performed immunofluorescent staining of nerve tissue. We found that dinutuximab has detectible binding to rat sympathetic nerve ganglion cells but INV721 does not (**Fig 6A**). Similarly, dinutuximab also shows detectable binding to nerve fibers of human peripheral nerves while INV721 does not (**Fig 6B**). By immunofluorescent staining, both dinutuximab and INV721 bind similarly to CHLA20 neuroblastoma xenograft tissue (**Fig 6A** and **6B**). By flow cytometry, we also found substantial binding of dinutuximab to nerve cells isolated from mouse dorsal root ganglion, while INV724 shows minimal binding over background (**Fig 6C, Supplemental Fig. 3**).

**Figure 6.**
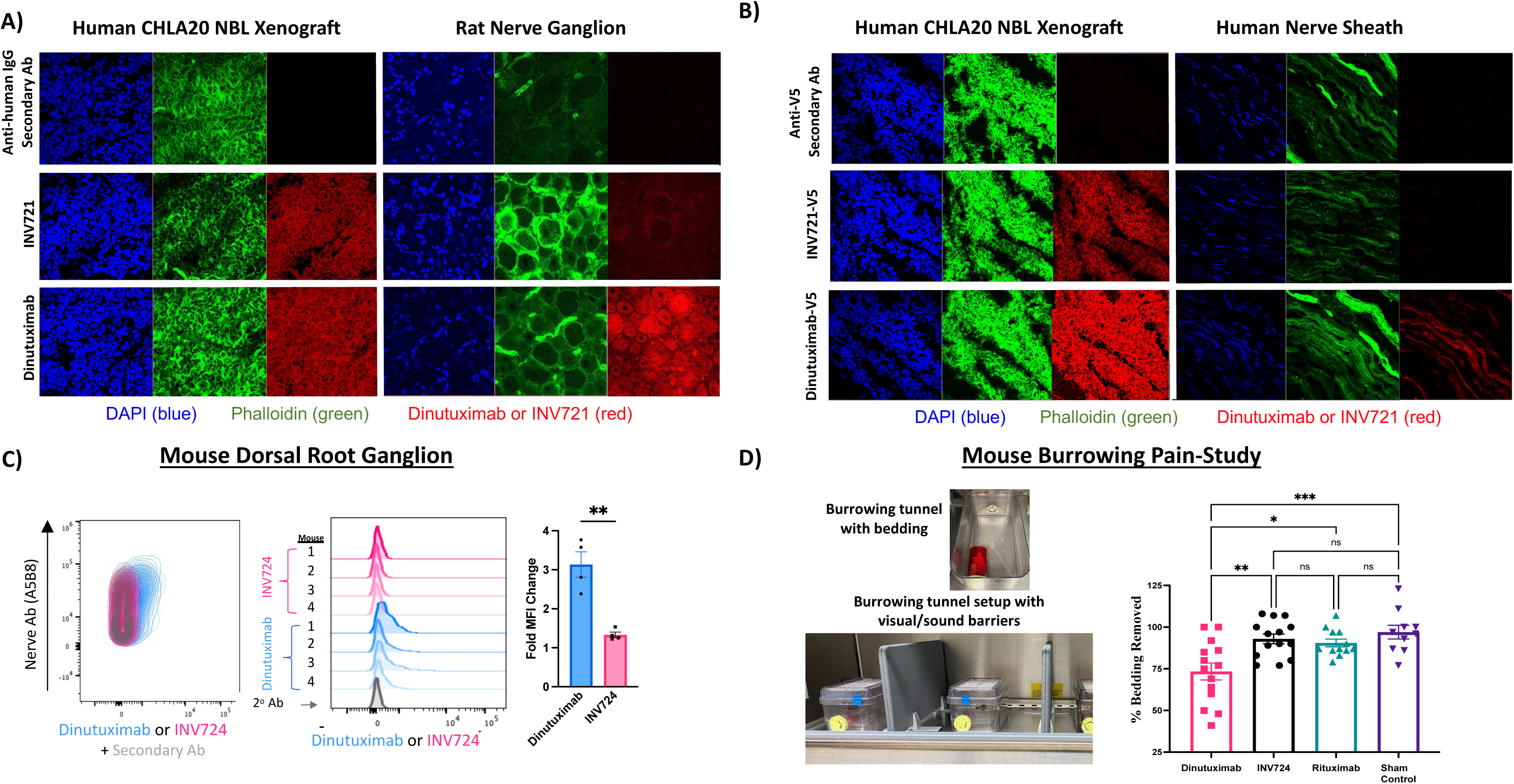
Anti-GD2 x B7-H3 does not bind nerves and does not cause pain in mice, whereas dinutuximab does. (A and B) Immunofluorescence analyses of human CHLA20 neuroblastoma xenografted tissue and rat nerve ganglion (**A**) or human peripheral nerve sheath (**B**) were labeled with dapi (blue), Phalloidin (green), as well as with INV721 (2^nd^ row) vs. dinutuximab (3^rd^ row), vs. no antitumor antibody (top row), followed by a secondary antibody (red) against human IgG (**A**) or against the V-5 tag on the antitumor antibodies (**B**). While CHLA20 human neuroblastoma xenograft tumors excised from NRG mice can be bound by both INV721 and dinutuximab (**A** and **B**), INV721 binds neither rat ganglion **(A)** nor human peripheral nerves (**B**). However, dinutuximab binds both rat ganglion **(A)** and human peripheral nerves **(B)**. Staining of slides was performed two separate times with similar results observed. **C)** Dorsal root ganglion nerve cells isolated from mouse spinal cord are bound by dinutuximab, but they are not bound by INV724 as assessed by flow cytometry (n=4 mice/group). A cell density plot (left panel) shows a shift in bound dinutuximab (blue) but not INV724 (pink). Histogram data for each individual mouse (middle panel) depicting increased bound secondary antibody to either dinutuximab-bound nerves (blue) as compared to INV724-bound nerves (pink) vs. secondary antibody alone (gray). The fold change over secondary alone (i.e., no primary antibody) of dorsal root nerve cells bound by either dinutuximab or INV724-showed a significant increase in the Mean Fluorescence Intensity (MFI) if incubated with dinutuximab as compared to INV724 (right panel). **D)** A burrowing study was performed to assess pain in C57Bl/6 mice. This assay measures the amount of bedding removed from tunnels placed in each cage from a single mouse (left upper image); cages with tunnels were separated by visual and sound barriers (left lower image). Mice were treated via IV injection with 250 µg of INV724, dinutuximab or rituximab, or they served as a no-treatment/needle-poked sham controls. The % change in burrowing activity (over 15 min with the burrowing tunnel) from time 0 (prior to antibody injection) to 1 hr-post antibody injection was calculated. Mice injected with INV724 removed a similar amount of bedding as the rituximab or non-handling group (no differences seen). In contrast, mice injected with dinutuximab removed significantly less bedding compared to all 3 other groups, including the INV724 group, indicating they were feeling pain, while the INV724 mice were not. n=10-14 mice/group, reflecting the combined results of 3 separate independent experiments.

Critically linked to binding of mAb to GD2 on nerve tissue is whether or not pain is experienced with treatment. Many animal models of pain using stimulus-evoked measures, such as the direct application of heat or pressure, have failed in the past to translate into clinical applicability.^41, 42^ Thus, we used a model of spontaneous pain, as spontaneous pain measures may be more relevant to the human condition of pain due to systemic injection of anti-GD2 antibody.^43^ The burrowing test measures housekeeping activities of mice that are essential for a satisfactory “quality of life” that correlate to normal daily human activities; reduced ability to do such activities are indicative of stress due to pain felt in mice.^44, 45, 46, 47^ Using this approach, as shown in **Fig. 6D** mice treated with dinutuximab show a significant decrease in displaced bedding compared to rituximab (control mAb) and sham control. In contrast, the amount of bedding displaced by INV724 treated mice was not different from the controls. These findings indicate that dinutuximab caused objectively measurable stress/pain, which was not caused by the INV724 or rituximab.

## Discussion

Creating this tumor-targeting bsAb that requires the co-expression of two separate tumor antigens (GD2 and B7-H3), improves the tumor-specificity over monospecific anti-GD2 or anti-B7-H3 mAbs. While B7-H3 is expressed on neuroblastoma, it is not found on nervous tissue. Here, we have shown that INV724 binds strongly to cells that express both GD2 and B7-H3, but it does not bind to nervous tissue (which expresses GD2 but not B7-H3), confirming that INV724 is more tumor-specific than dinutuximab or other anti-GD2 mAbs. Separately, we showed similar antitumor efficacy between dinutuximab and INV724 in *in vitro* ADCC assays and in *in vivo* GD2^+^/B7-H3^+^ tumor models. Finally, we confirmed that while dinutuximab binds nerve cells [ganglion (rat), myelin fibers (human) and dorsal root ganglion (mouse)], INV724 does not. Corresponding to these findings, mice do not display symptoms of neuropathic pain if treated with INV724, but they do show symptoms of neuropathic pain when treated with dinutuximab. These preclinical data suggest that clinical administration of INV724 to patients with neuroblastoma might enable similar clinical benefit, with a substantial decrease in neuropathic toxicity in settings where anti-GD2 mAbs are currently being used. Clinical Phase-I and Phase-II testing will be needed to determine this.

The current maintenance regimen for patients receiving dinutuximab involves 4 consecutive daily 4-20 hour IV infusions; if this is tolerated, they then receive 4 more 4-day courses, every 3-4 weeks. Yet, the administration of dinutuximab is associated with a high-degree of toxicity, including, grade 3 or 4 neuropathic pain reported in 52% of patients.^1, 3^ In many cases, this intense pain prevents patients from receiving the entire full-dose dinutuximab treatment regimen. The type of pain reported by neuroblastoma patients receiving anti-GD2 mAbs includes generalized pain, as well as abdominal, extremity or back pain, neuralgia (stabbing/burning feeling) and arthralgia (joint pain).^1, 48^ Therefore, nearly all patients treated with dinutuximab (and other anti-GD2 mAbs) require opioid treatment given concurrently, and many require other agents, such as IV lidocaine, or sedation, to alleviate the acute painful side-effects. By enhancing tumor-specificity with INV724, and eliminating binding to nervous tissue, we should substantially reduce the painful neuropathic side-effects that patients experience during anti-GD2 administration. By reducing, or eliminating, the painful side effects associated with anti-GD2 immunotherapy, it may be possible to avoid the use of opioids and other IV analgesics.^49^ Less infusion associated neuropathic pain may also allow for outpatient antibody administration and improved quality of life during the multiple courses of antibody treatment. In addition, the reduced pain may allow for increased tolerability, with more patients able to receive the full dose and scheduled course of therapy, potentially leading to improved outcomes.^50^

Prior to the development of this GD2xB7-H3 bsAb, other efforts to decrease the neuropathic pain associated with anti-GD2 mAb have been reported. Substantial work has been done with the hu14.18K322A mAb which has a point mutation in the IgG heavy chain to decrease complement activation. As complement activation has been shown to be at least part of the mechanism for neuropathic pain following anti-GD2 mAb treatment, this agent has been used in attempt to decrease this toxicity.^10, 12, 13, 51^ Patients receiving hu14.18K322A mAb do appear to experience less pain than patients receiving dinutuximab, with an MTD of 60mg/m^2^/dose (2-3 times higher than the MTD of dinutuximab given as single agent). Yet, patients receiving hu14.18K322A mAb still experience neuropathic pain, accounting in part for its MTD.^12, 13, 51^ Recently, a version of the ch14.18 anti-GD2 mAb with an IgA Fc was developed that can cause ADCC (particularly by neutrophils), while not activating complement via C1q or inducing neuropathic pain (as tested by the Von Frey method). Clinical testing of this mAb has not yet begun.^52^ In parallel, a separate anti-GD2 mAb has been used with anti-CD3 antibody as a bifunctional T cell engager, using an anti-GD2 component that does not activate complement. It is potent *in vitro* and in mice, but clinical testing has not been reported.^53^ Separate work has indicated that while neuroblastoma and nerves both express GD2, neuroblastoma, but not nerves, has a modified O-acetyl-GD2.^39, 54, 55^ A mAb against the O-Acetyl-GD2 has been developed, that like INV724, binds to neuroblastoma but not to nerves. Preclinical testing in mice with the anti-O-Acetyl-GD2 mAb suggests it may have antitumor activity without inducing pain. Clinical testing has not been reported. To what degree neurotoxicity and pain are caused by complement remains uncertain, since patients treated with the hu14.18K322A still do have pain, but to a lesser degree than with treatment with other anti-GD2 mAbs currently in use.^12^ It does seem that the neuropathic pain induced by these anti-GD2 antibodies, compared to the many other antitumor mAbs approved or in clinical testing that do not bind to nerves and do not cause neuropathic pain (such as rituximab or cetuximab), relates to the unique ability of the anti-GD2 mAbs to bind to peripheral nerves.^14, 15, 38, 56, 57^ Thus, since therapeutics that bind monospecifically to GD2 can also bind to nerves, it seems possible that potent forms of any anti-GD2 therapy may cause neuropathic pain and potentially risk nerve damage. One way to potentially avoid this is to use a therapeutic that does not bind monospecifically to GD2, and thus cannot bind to nerves (as shown here for INV724).

Besides developing modifications to anti-GD2 Abs to reduce toxicity, another approach to targeting neuroblastoma is to target different surface expressed tumor-antigens not often found on normal tissue. One such antigen is B7-H3. Ongoing clinical testing of mAbs targeting B7-H3 are being tested for neuroblastoma treatment. As the major mechanism of Ab-based immunotherapy in neuroblastoma is via inducing ADCC or antibody dependent cellular phagocytosis (ADCP), we compared a bivalent B7-H3 targeted mAb, I7-01, which has high avidity to B7-H3 similar to anti-B7-H3 antibodies being used in clinical testing for neuroblastoma. We observed greater ADCC potential against neuroblastoma targets using INV724 or anti-GD2 mAbs as compared to an anti-B7-H3 mAb (**Fig. 3**). We hypothesize that this enhanced ADCC may be due to two separate factors: 1) the far greater expression of GD2 than B7-H3 on most human neuroblastomas, and 2) the proximity of the GD2 antigen to the surface of the cells, whereby the GD2 antigen draws the bound antibody much closer to the cell surface than an anti-B7-H3 mAb does due to the size of the B7-H3 protein (**Supplemental Fig. 4**). As such, monospecific/bivalent conventional anti-B7-H3 mAbs may have difficulty achieving high levels of binding to cells that have low, but detectable, levels of B7-H3. IIn contrast, once the anti-B7H3 Fab of an INV724 molecule has bound its B7-H3 target, the anti-GD2 Fab should rapidly find a highly expressed, membrane-embedded, GD2 target, enabling bivalent high avidity binding and thus resulting in more INV724 binding and ADCC efficacy (**Figs. 2A&3D**).

Recent studies indicate that anti-GD2 mAbs can be more effective when given with chemotherapy.^58^ Based on these findings and other current insights, anti-GD2 antibody has shown more rapid partial and complete responses when added to induction chemotherapy for newly diagnosed neuroblastoma patients at St. Jude.^10, 59^ Based on this potential benefit for anti-GD2 antibody during the induction chemotherapy and the efficacy shown for relapsed patients when anti-GD2 is given with chemotherapy, the Children’s Oncology Group is moving to a randomized trial that includes dinutuximab in maintenance and randomizes patients to induction with or without anti-GD2 [NCT06172296].^58, 59^ This potential increased applicability of anti-GD2 therapy, for use in induction, maintenance and relapse, makes this work with INV724 potentially more applicable in these clinical settings, if INV724 proves clinically to decrease or avoid the pain of treatment.

Finally, if the pain is significantly reduced, it may be possible to tolerate substantially higher doses of INV724 than is currently used for dinutuximab (or other anti-GD2 mAbs, which have dosing limited by pain). The current dose of dinutuximab per treatment course is only 5-10% of the current dosing used per course for other antitumor mAbs that do not cause neuropathic pain (such as the anti-CD20 mAb, rituximab; or the anti-EGFR mAb, cetuximab).^5, 6^ Our lab has shown (by analyses of anti-GD2 plasma levels in neuroblastoma patients receiving anti-GD2 mAb) in three separate clinical trials that patients with higher plasma levels of anti-GD2 mAb show better clinical antitumor outcome.^10, 17, 58^ Based on this, we hypothesize that our ability to give meaningfully higher doses of INV724, and/or an extension to the number of courses, than is currently used for dinutuximab (closer to the dosing used for rituximab and cetuximab) may translate to an improved clinical antitumor effect than is currently obtained with dinutuximab or other anti-GD2 mAbs. Clinical phase-I testing will be needed to determine the *in vivo* clinical toxicity of INV724, and its associated maximal tolerated dose.

In summary, these preclinical data support initiating clinical phase-I testing of this GD2xB7-H3 baAb. If it is tolerated at higher doses or for longer periods of time than current anti-GD2 mAbs, it may potentially be tested for efficacy in the same clinical settings; namely with chemotherapy during induction, with cis-retinoic acid (and GM-CSF) for maintenance, and with chemotherapy for relapsed disease. While clinical testing and development will be required, our goal is to enable better antitumor efficacy with less neuropathic pain.

## Online Methods

### Methods

#### Construction of B-Body Bispecific Antibodies to B7-H3 and GD2

Bispecific antibodies (bsAbs) were created from B7-H3 and GD2 antibodies as depicted in Supplemental Figure 1 by reformatting the B7-H3 antibodies into the first and second polypeptide chains and GD2 into the third and fourth polypeptide chains in an antibody construct. The four polypeptide chains were transiently transfected into HEK cells to produce the antibodies. The bispecific antibodies secreted into the cell culture medium were subsequently isolated with an anti-CH1 affinity capture resin followed by polishing using a strong cation exchange resin. Bispecific constructs in which either the B7-H3 or the GD2 arm was replaced with a non-binding arms were also created.

#### Production of INV721 and INV724 (Afucosylated version)

INV721 and INV724 were produced from CHO cell lines stably expressing the molecules. For the afucosylated version (INV724), the expresson of alpha-(1,6)-fucosyltransferase is inhibited.

#### Cell Lines

Human neuroblastoma cell lines (LA-N-1, LA-N-5, NGP, NB69, NB-EBc1, IMR5, NLF and SK-NAS) were kindly obtained from Dr. John Maris of the Children’s Hospital of Philadelphia, and were grown in RPMI 1640 supplemented with 10% heat-inactivated FBS, 2 mmol/L l-glutamine, 100 U/mL penicillin, and 100 µg/mL streptomycin. The human neuroblastoma line (CHLA20) was grown in DMEM supplemented with 10% heat-inactivated FBS, 2 mmol/L l-glutamine, 100 U/mL penicillin, and 100 µg/mL streptomycin.

The murine B78-D14 (B78) melanoma is a poorly immunogenic cell line derived from B78-H1 cells, which were originally derived from B16 melanoma and kindly obtained from Ralph Reisfeld, PhD, The Scripps Research Institute, La Jolla, CA.^60, 61, 62^ B78-D14 cells have functional GD2/GD3 synthase and express the disialoganglioside GD2.^60, 61^ B78 cells were grown in RPMI 1640 supplemented with 10% heat-inactivated FBS, 2 mmol/L l-glutamine, 100 U/mL penicillin, and 100 µg/mL streptomycin. Periodic treatment with Hygromycin B (50 _μ_g/mL) and G418 (400 _μ_g/mL) was used to maintain GD2 expression of B78-D14 cells.

The NXS2 cell line (kindly obtained from Ralph Reisfeld, PhD, The Scripps Research Institute, La Jolla, CA, and then maintained by Alice Yu, MD, University of California, San Diego, CA) is a moderately immunogenic, highly metastatic, GD2-positive murine neuroblastoma cell line.^63^ NXS2 is a hybrid between GD2-negative C1300 (a neuroblastoma tumor that spontaneously arose in A/J mice^64^) and GD2-positive murine dorsal root ganglion cells (C57Bl/6 J background), but does not express C57Bl/6 H-2 and therefore grows in immunocompetent A/J mice. NXS2 cells were grown in DMEM medium supplemented with 10% FBS, 2 mM L-glutamine, and 100 U/ml penicillin/streptomycin.

All cell lines used were routinely monitored for mycoplasma by PCR testing as previously described.^65^

#### Cell Line Modifications

B78-D14s were modified to knock-out expression of murine B7-H3 via CRISPR knock-down. In brief, in a 96 well plate, B78 cells at 50% confluency were treated with TrueGuide™ Synthetic sgRNA targeting B7-H3 (ThermoFisher, cat. no. CRISPR142889_SGM) using TrueCut^™^ Cas9 Protein v2 (ThermoFisher, cat. no. A36496) and Lipofectamine^™^ CRISPRMAX^™^ Cas9 Transfection reagent (ThermoFisher, cat. no. CMAX00001) following the manufacturers protocol. Cells were transferred to a 6 well plate, single-cell cloned, and assessed for expression of mouse B7-H3 by flow cytometry compared to B78-D14 cells not treated with sgRNA targeting B7-H3. For testing by flow cytometry, cells were labeled with anti-mouse B7-H3-APC antibody (Biolegend, cat. no. 135608) for 30 min, washed and analyzed on an Attune NxT Flow Cytometer (ThermoFisher). B78-D14 clones that showed 50% decrease in murine B7-H3 were treated again with sgRNA targeting B7-H3 following the manufacturers protocol, followed by single-cell cloning and assessment by flow cytometry for murine B7-H3 expression. A B78-D14 clone that had lost expression of murine B7-H3 (B78-mB7-H3) was selected following two treatments with B7-H3-sgRNA treatment.

After confirmating that B78-mB7-H3 cells did not express murine B7-H3 by flow cytometry, human B7-H3 was transduced into B78-mB7-H3 and NXS2 (that still had mB7-H3) by lentiviral transduction using pLV-mCherry:T2A:Puro-EFS>hCD276 [NM_001024736.1] (VectorBuilder; Vector ID VB170825-1084zey). Positively transduced cells were selected for using puromycin (4 µg/ml for B78, 2 µg/ml for NXS2). For B78 cells, stably transduced cells, referred to as B78-mB7-H3+hB7-H3, were then single-cell cloned and tested for GD2 and human B7-H3 expression. Separate B78-mB7-H3+hB7-H3 clones were generated, including B78-mB7-H3+hB7-H3 Cl.6 (GD2+/hB7-H3+; used for *in vivo* efficacy testing). Separate clones were expanded based their variable expression of GD2 and/or B7-H3 and used for *in vitro* specificity, *in vitro* efficacy/ADCC and PET imaging studies, including B78-mB7-H3+hB7-H3 Cl. 8 (GD2-/hB7-H3-); B78-mB7-H3+hB7-H3 Cl. 13 (GD2+/hB7-H3+); B78-mB7-H3+hB7-H3 Cl. 14 (GD2-/hB7-H3+); B78-mB7-H3+hB7-H3 Cl. 25 (GD2+/hB7-H3-). For NXS2, stably transduced cells, referred to as NXS2+hB7-H3, were sorted, single-cell cloned and tested for GD2 and human B7-H3 expression.

For *in vivo* IVIS imaging, CHLA20 cells were transduced as above with lentiviral transduction using pLV-Puro-EFS>TurboGFP:IRES:Luciferase (VectorBuilder; Vector ID VB180725-1103sza). Positively transduced cells were selected for using puromycin (2µg/ml).

#### Cell Binding

B7-H3xGD2 and the corresponding B7-H3xNon-Binding arm or GD2xNon-Binding arm bsAbs were tested by flow cytometry for cell binding on a human B7-H3-expressing B78 murine melanoma cell lines which overexpressed both human B7-H3 and GD2 or only GD2. Cells were incubated for 60 minutes at 4°C with a dilution series of the bsAbs ranging from 333 nM to 0.02 nM prepared in PBS (Corning, #21-031-CV). Cells were then washed with PBS and resuspended in a secondary antibody solution of AF488 goat anti-human IgG Fab Fragment (Jackson Immuno Research, #109-547-003) diluted in PBS. Cells were incubated in the secondary antibody solution for 30 minutes at 4°C, washed in PBS, resuspended in cold PBS, and subjected to flow cytometry to determine the MFI.

#### Measurement of Binding Affinities

Monovalent affnities to B7-H3 and GD2 were determined by biolayer interferometery (BLI) on the OctetQK384 system (Pall ForteBio). Biotinylated B7-H3 or GD2 was immobilized on streptavidin sensors. The antigen-immobilized sensor was submerged into a solution containing various concentrations of bispecific antibody ranging from 3-200 nM for B7-H3 or 150 nM – 10 _μ_M for GD2. The real-time association and dissociation curves of the bispecific molecule were fit using ForteBio software in a 1:1 binding model with global or local fit to obtain association (kon) and dissociation (koff) rates. *K*_d_ was calculated from these rate constants.

#### Antibody Dependent Cellular Cytotoxicity

Target tumor cell lines transduced to express nuclear localized mKate2 (NLS-mKate2) were plated at 100 cells/well or 250 cells/well (depending on cell size) in 20 µL/well in a 384-well round bottom spheroid plate (Corning, #4516). Cells were incubated at 37 °C for 48 hrs to allow spheroids to form. During that incubation, peripheral blood mononuclear cells (PBMCs) were isolated from healthy donors as previously described and incubated overnight at 37 °C in RPMI 1640 supplemented with 10% heat-inactivated FBS, 2 mmol/L l-glutamine and 200 U/ml IL-2.^4^ PBMCs were centrifuged at 350 x g for 10 minutes, washed with 50ml of PBS to remove IL-2, and resuspended in RPMI 1640 supplemented with 10% heat-inactivated FBS, 2 mmol/L l-glutamine. PBMCs were added to spheroids in a 384-well plate in 20 µL of media/well at a ratio of 2,500 PBMCs to 1 spheroid. Antibodies were added in 10 µL of media/well, with the final concentration of antibody added dependent upon the experiment. Staurosporine was added to cause maximum tumor death at a 1mM/well. The final volume of all wells was brought to 50 µL/well with media. Plates were placed in an Incucyte S3 system (Sartorius), and imaged every 4 hrs using the Spheroid Module.

#### FC_Y_RGenotyping Internalization Assays

Genotyping for FC_Y_R3A, FC_Y_R2A and FC_Y_R2C and categorical determinations for high-affinity FC_Y_Rs vs low-affinity FC_Y_Rs were performed as previously described.^66, 67, 68^

#### Antibody Internalization Assays

Antibodies were labeled with pHrodo^TM^ iFL Red (ThermoFisher) per the manufacturer’s protocol. Antibodies were plated together with neuroblastoma cell lines and imaged every 3-6 hours for 48 hours via an Incucyte S3 live cell imaging platform.

#### Animal Models

Female 6-8 week-old C57BL/6 (Taconic Farms; strain B6NTac) and A/J mice (Jackson Labs; strain Strain #:000646) were used for these studies involving syngeneic tumor models (B78 and NXS2, respectively). Mice were housed in the University of Wisconsin-Madison animal facilities at the Wisconsin Institutes for Medical Research. Mice were used in accordance with the *Guide for Care and Use of Laboratory Animals* (NIH publication 86-23, National Institutes of Health, Bethesda, MD, 1985).

Intradermal (i.d.) tumors were established by injecting 2×10^6^ B78-mB7-H3+hB7-H3 Cl.6, or NXS2+hB7-H3 in Matrigel (Sigma, # 354248), in 0.1 ml of PBS into the shaved flank of mice.^69^ Tumor diameters were measured, and tumor volume (mm^3^) was calculated as ½ x tumor length x tumor width^2^. One day before RT, mice were randomized according to tumor volumes and assigned to experimental groups. Naïve mice were injected with the same tumor cells as a control.

Female 6-8 week-old nude mice (Strain #:007850 RRID:IMSR_JAX:007850) were used for studies involving human neuroblastoma tumors (CHLA20-luciferase). Mice were housed in the University of Wisconsin-Madison animal facilities at the Wisconsin Institutes for Medical Research. Mice were used in accordance with the *Guide for Care and Use of Laboratory Animals* (NIH publication 86-23, National Institutes of Health, Bethesda, MD, 1985).

Intravenous (IV) metastatic tumors were established by injecting 5×10^5^ CHLA20-luciferase in 0.1 ml of PBS into the tail vein of mice. Tumor development was monitored weekly by bioluminescent imaging with Perkin Elmer IVIS Spectrum In Vivo Imaging System and Living Image Software (Caliper Life Sciences, Hopkinton, MA) was used for image processing. Bioluminescence was measured 15 minutes following intraperitoneal injection of 200 µL (150 mg/kg) of luciferin.

#### Burrowing Method to Monitor Pain

An assay for testing mouse burrowing behavior was developed based on previously published reports.^44, 46, 47, 70^ Mouse burrowing apparatuses were constructed in house. A red mouse tunnel (VWR, cat no. K322) was fit with a plastic circle that was cut to size to adhere to one end of the tunnel with tape. Tubes were labeled A-D. Mice housed in groups of 5 mice/cage were acclimated to an empty burrowing apparatus on Day 0 by placing the tunnel in each cage of the mice (labeled A-D) for 4 hrs. The following day, burrowing activity was tested. Empty test-cages (labeled A-D) were separated with noise/visual barriers within a biosafety cabinet with the blower turned on **(Fig 5C)**. To test for burrowing activity, individual mice were removed from their home cage and placed in an empty new cage with their corresponding home-cage tunnel (e.g. cage labeled “A” with tunnel labeled “A”) filled with 30g of bedding **(Fig 5C)**. The bedding that was used included 25g of fresh bedding from an unused mouse cage combined with 5g of bedding from the home-cage of the mice. Bedding was stored and saved in plastic bags (labeled A-D) for the remaining days of the study.

On Day 1 at 9am, mice were tested for pre-burrowing to ensure they would burrow. Mice were left alone for 15 min, and the amount of bedding removed from the burrow was measured. Mice were then returned to their home cage. On Day 2 at 9am, mice were tested for pre-treatment burrowing at time 0 as described above. Mice that removed >20g of bedding were enrolled in the study.

After the pre-treatment burrowing evaluation, enrolled mice were restrained in a plastic holder and injected via tail vein with 200µL of their treatments. Treatment groups included dinutuximab (250µg/dose), INV724 (250µg/dose), rituximab (250µg/dose), or a needle-poke control. Each cage included mice with different treatment groups (e.g. cage A had mice treated with dinutuximab, INV724, rituximab and needle-poke control) to control for emotional contagion. One hour after injection with Ab, mice burrowing activity was tested again. Experimenters assessing burrowing behaviors were blinded to which treatment mice received.

The % change in burrowing activity from Day 2 time pre-burrow to Day 2 time 1 hr-post treatment was calculated. Burrowing studies were repeated 3 separate times, with 3-5 mice enrolled per treatment group during each study for a final total of 10-14 mice/treatment group.

#### PET Imaging of Tumors

NRG mice were injected with 10^6^ tumor cells intradermally^69^ with NGP tumor cells on the lower right flank. For the 4 tumor model, C57Bl/6 mice were injected with 10^6^ tumor cells intradermally^69^ with B78-mB7-H3+hB7-H3 Cl. 8 (GD2-/hB7-H3-) on the upper right flank; B78-mB7-H3+hB7-H3 Cl. 13 (GD2+/hB7-H3+) on the lower right flank; B78-mB7-H3+hB7-H3 Cl. 14 (GD2-/hB7-H3+) on the upper left flank; and B78-mB7-H3+hB7-H3 Cl. 25 (GD2+/hB7-H3-) on the lower left flank (**Fig. 4D**). When the tumors reached ∼100-300 mm^3^, the mice were injected with Zr^89^-labeled antibodies prepared as described previously.^71^ Briefly, p-SCN-Bn-Deferoxamine (Df) was conjugated by thiourea linkage to the antibody at pH ∼8.5 and purified using size exclusion chromatography. Conjugated antibody was radiolabeled with ^89^Zr produced on a GE PETtrace cyclotron by using the ^89^Y(p,n)^89^Zr reaction. For radiolabeling, ^89^Zr-oxalate was added to the chelator conjugated antibody and incubated in 1 M HEPES buffer (pH=7.5) at 37°C for 1 h at a ratio of 100 _μ_g antibody per mCi of ^89^Zr. Radiolabeled antibody (^89^Zr-Df-INV724) was purified using size exclusion chromatography. Mice were intravenously injected with 5.55 – 9.25 MBq (150-250 _μ_Ci) for PET imaging studies. PET/CT images were decay corrected and images are normalized to percent injected dose per gram of tissue (%ID/g) using the Inveon Research Workspace by manually drawing volumes-of-interest.

#### In Vivo Tumor Model Efficacy Studies

Mice bearing GD2/B7-H3-expressing melanoma (B78) or neuroblastoma (NXS2) tumors were treated with our *in situ* vaccine regimen: external beam radiation therapy (RT, 12Gy, Day 1) followed by intratumoral injection of IL2 (75K U/dose) with or without INV724 (40µg/dose, Day 5-9) or dinutuximab (40µg/dose, Day 5-9). Mice were monitored for tumor growth via (twice weekly caliper measurements). Mice were euthanized when either the length or the width of a tumor reached 20mm and tracked for survival.

Nude mice were injected via tail vein with CHLA20-luciferase tumors on Day 0, followed by retro-orbital injection of INV724 (25µg/dose) or dinutuximab (25µg/dose) in 0.1ml of PBS, or PBS alone, on D1, D4, D7 and D10. Retro-orbital injections were performed using sterile technique under isoflurane anesthesia.

#### Antibody Titrations

Tumor cells were harvested and resuspended at 5×10^6^ cells/ml in flow buffer. From this suspension, 50µL of cells were added to flow cytometry tubes, for a total of 250,000 cells/tube. Serial dilutions of primary antibody (INV724 or dinutuximab) were created, and 50µL of each antibody dilution was added its respective tube. Cells were incubated for 30 min in the dark at 4C and washed with flow buffer (PBS + 2% FBS). A secondary master mix was made by diluting anti-Fab-PE (109-117-008; Jackson ImmunoResearch labs) 1:50 in flow buffer, and 100µL of the diluted secondary antibody was added to the tubes and incubated for 20-30 minutes in the dark at 4C. Samples were washed with flow buffer, DAPI was added, and cells were subjected to flow cytometry on an Attune NxT flow cytometer to determine the MFI.

#### Antibody Density Testing

Cells were harvested and added to flow tubes (200,000/100µl flow buffer) for staining with GD2 APC or CD276 APC (BioLegend, 357306 and 351006). Quantum Simply Cellular anti-mouse IgG beads (Bangs Laboratories, 815A) were prepared in flow buffer in microcentrifuge tubes and stained at the same time. Amounts of antibody added to the beads and to each cell line were determined previously by antibody saturation titration (additional 50% antibody volume added until less than a 10% increase in fluorescence is obtained=antibody saturation). Cells and beads were incubated for 30 min in the dark at 4°C, and washed with flow buffer. Beads were washed an additional two times as per manufacturer’s recommendations. DAPI was added to cell samples before running on an Attune NxT flow cytometer and beads were collected after the cells at the same instrument settings. Bangs QuickCal template for BD Relative Linear Channels (1 - 10,000 channels, for Log Data) and bar (range) gates for the bead populations (about 1000 beads per tube collected) were used for Antigen Binding Capacity (ABC) analysis and the unstained control ABC result was subtracted from the test ABC result for each cell line. If monovalent antibody to receptor binding is presumed, the ABC value is the number of surface receptors.

#### Immunofluorescent Staining of Tissue

Rat peripheral nerve tissue and CHLA20 tumors grown in nude mice were harvested, embedded in OCT without fixation, and frozen on liquid nitrogen. Embedded tissues were cryosectioned at 10 micron sections onto Colorview adhesion slides (StatLab). OCT-embedded human peripheral nerve tissue (specimen 407E) was a kind gift from Ambsio and sectioned at 7 microns per tissue onto Superfrost slides (ThermoFisher). Following a slow thaw after removal from -80C storage, slides were incubated in cold acetone (maintained at -20C) for 10 min and then dried for 10 min at RT. Slides were rinsed under water for 10 min, rinsed with PBS 3 times, and blocked with 10% FBS for 1 hr at RT. For rat nerve tissue, primary INV724 (Invenra) or dinutuximab antibodies (commercial grade, Unitixun) at 3 µg/ml in 1% FBS, or unstained sections in 1% FBS block, and incubated overnight at 4C. Slides were washed 3 times in PBS, and 1%FBS with 1 µg/ml of secondary anti-human-IgG-AF555 (A-21433, LifeTech) was added and incubated for 1 hr at RT. For human peripheral nerve staining, sections were stained with primary INV724+V5 tag or dinutuximab+V5 tag antibodies (Invenra) at 3 µg/ml in 1% FBS, or unstained sections in 1% FBS block, were incubated overnight at 4C. Slides were washed 3 times in PBS, and 1%FBS with 1 µg/ml of secondary anti-V5-AF55 (49355S, Cell Signaling) was added and incubated for 1 hr at RT. Slides were washed 3 times with PBS, and fixed in 10% neutral buffered formalin for 10 min. Slides were washed 3 times in PBS, and incubated with phalloidin-AF488 (15nM, A12379, ThermoFisher) for 30 min at RT. Slides were washed with PBS 3 times, with a drop of DAPI being added with the third wash. Slides were coverslipped with Prolong Gold mounting media (ThermoFisher) and imaged on a confocal microscope (LSM 710). These staining procedures were repeated twice with similar results.

#### Dorsal Root Ganglion Flow

Dorsal root ganglion were isolated from 4 naïve C57Bl/6 mice and digested into single cell suspensions.^72^ Single cell suspensions were stained with 1µg of dinutuximab or INV724 and incubated at room temperature for 20 min. Cells were washed with flow buffer, centrifuged (350g X 5 min), and incubated with secondary anti-human IgG1-PE (MA1-10389, ThermoFisher) for 20 min room temperature. Cells were washed with flow buffer, centrifuged, and incubated with A2B5-AF657 antibody (to stain dorsal root ganglion nerve cells) for 20 min.

Cells were washed, centrifuged, and DAPI was added prior to flow cytometry analysis on an Attune NXT (ThermoFisher). Data were analyzed using FlowJo software by gating on live, A2B5+ nerve cells, and assessed for the amount of bound secondary PE antibody by Median Fluorescence Intensity (MFI) to either dinutuximab or INV724. This study was repeated twice with similar results.

#### Statistics

Tumor spheroid shrinkage data were compared by one-way ANOVA followed by Tukey’s Post Hoc test for multiple comparisons. For statistical analysis of tumor growth curves, the time-weighted average (area under the volume-time curve, calculated using trapezoidal method) was calculated for each mouse tumor. Time-weighted averages were compared between treatment groups overall by a Kruskal-Wallis test, and since significance was reached, then pairwise by Mann-Whitney-Wilcoxon tests. Survival curves were compared with pairwise log rank tests and response was evaluated by two-sample tests of proportions. Chi-square test was used to assess differences in response rate (i.e., tumor free vs. tumor bearing). Significance was assessed at the alpha = 0.05 level and no adjustments were made to account for inflated type 1 error rate. Analysis was conducted using R version 4.3.1 (2023-06-16).

## Supplemental Materials

**Supplementary Figure 1: FC_Y_R genotypes influence ADCC activity.** ADCC was performed using PBMCs from 6 different donors using 40ng/ml of each antibody tested. Donors with medium to high-affinity FC_Y_R genotypes (top panel) showed similar ADCC capabilities by INV721 and INV724, regardless of fucosylation status. Donors with low-affinity FC_Y_R genotypes (bottom panel) showed the strongest ADCC when using the afucosylated INV724 antibody (INV724). For each of the 6 donors, the specific amino acid genotype at the critical allele location in the FC_Y_R 3A (V or F) and FC_Y_R 2C (C or T) that determinies their high or low affinity genotypes, as previously published, is indicated.^32, 33, 67, 68^

**Supplementary Figure 2: Antigen density of B7-H3 and GD2 on melanoma cell lines.** The antigen densities (surface receptors per cell) of GD2 (white bars) and B7-H3 (laddered black lined bars) were determined for murine B78 melanoma cell line GD2/B7-H3 variants transduced to express human B7-H3, B78 cells not transduced to express human B7-H3, murine B16 melanoma (which does not express GD2 or human B7-H3), and human M21 melanoma. As B16 cells do not express either GD2 or human B7-H3, they were used as the standard in this assay to set a cut-off for detectible antigen expression (as indicated by the dashed horizontal line at ∼10^3^ surface receptors per cell). Human M21 melanoma cells have endogenously expressed antigen densities of B7-H3 and GD2 that are similar to those expressed by the B78-Variant 4 (GD2^+/^B7-H3^+^).

**Supplementary Figure 3: Gating strategy for dorsal root ganglion nerves. (1)** Dorsal root ganglion cells were gated on by Forward Scatter Area (FSA) and by Side Scatter Area (SSA). Single cells were then gated on by selecting gates within FSA x FS-Height (FSH) (**2**) and then by SSA x SS-Height (SSH) (**3**), followed by gating on live DAPI-negative cells (**4**). The left plot of A2B5-AF647 Ab x Secondary PE Ab (**5**) shows that based on the no antibody control and the secondary PE Ab only control (no nerve A2B5-AF647 antibody) samples, gates for nerve positive samples were set. Cells within this gate labeled “Nerves with histo” (short for histogram) were used to create the histogram and cell contour plots displayed within Figure 6. The results are shown from 4 mice per group (individual mice labeled as DIN 1-4 or INV 1-4).

**Supplementary Figure 4: Antigen proximity and density influence ADCC potential.** On tumors from neuroectodermal origin, GD2 antigen (blue triangle) is expressed at a high density and can be targeted with an anti-GD2-mAb. Since the GD2 antigen is found at a high antigen density, anti-GD2 mAbs (pictured with 2 **blue** Fabs) can bind with high avidity and have high levels of tumor binding. B7-H3 (red rectangle), is expressed on multiple tumor types, including neuroblastoma, but is often found expressed at a lower density than GD2 (about 10-fold lower). B7-H3 can be targeted with an anti-B7-H3 mAb (pictured with 2 **red** Fabs), but because it is expressed at such low density on many tumors, the anti-B7-H3 mAbs can’t bind at high levels and thus less mAb remains bound for engaging immune functional activities. Using a B7-H3xGD2 bispecific antibody that can target B7-H3 with one Fab (**red**) and GD2 with the other Fab (**blue**), INV724 has preferential targeting to tumor cells, and can bind with high avidity to the target antigens, and more antibody is able to bind to the tumor, than for the B7-H3 antibody at low concentrations (see **Fig. 2A**), thus accounting for the greater killing by INV721 and INV724 than the anti B7-H3 mAb in **Fig. 3D**.

## Acknowledgements

This work was supported by Invenra Inc, The Wisconsin Alumni Research Foundation, Midwest Athletes Against Childhood Cancer, the University of Wisconsin Carbone Cancer Center and research grants from the Pablove Foundation, the HESI-thrive Foundation, the Hyundai Hope on Wheels Foundation, the End Kids Cancer Foundation and by public health service grants R35-CA197078, and P01 CA250972 from the National Cancer Institute.

## References

1. Armideo, E., Callahan, C. & Madonia, L. Immunotherapy for High-Risk Neuroblastoma: Management of Side Effects and Complications. J Adv Pract Oncol 8, 44–55 (2017).

2. Keyel, M.E. & Reynolds, C.P. Spotlight on dinutuximab in the treatment of high-risk neuroblastoma: development and place in therapy. Biologics 13, 1–12 (2019).

3. Yu, A.L. et al. Anti-GD2 antibody with GM-CSF, interleukin-2, and isotretinoin for neuroblastoma. N Engl J Med 363, 1324–1334 (2010).

4. Hank, J.A. et al. Augmentation of antibody dependent cell mediated cytotoxicity following in vivo therapy with recombinant interleukin 2. Cancer Res 50, 5234–5239 (1990).

5. Chang, M., Samlowski, W. & Meoz, R. Effectiveness and toxicity of cetuximab with concurrent RT in locally advanced cutaneous squamous cell skin cancer: a case series. Oncotarget 14, 709–718 (2023).

6. Held, G. et al. Radiation and Dose-densification of R-CHOP in Primary Mediastinal B-cell Lymphoma: Subgroup Analysis of the UNFOLDER Trial. Hemasphere 7, e917 (2023).

7. Erbe, A.K. et al. Neuroblastoma Patients’ KIR and KIR-Ligand Genotypes Influence Clinical Outcome for Dinutuximab-based Immunotherapy: A Report from the Children’s Oncology Group. Clin Cancer Res 24, 189–196 (2018).

8. Yu, A.L. et al. Long-Term Follow-up of a Phase III Study of ch14.18 (Dinutuximab) + Cytokine Immunotherapy in Children with High-Risk Neuroblastoma: COG Study ANBL0032. Clin Cancer Res (2021).

9. Cheung, I.Y. et al. Phase I trial of anti-GD2 monoclonal antibody hu3F8 plus GM-CSF: Impact of body weight, immunogenicity and anti-GD2 response on pharmacokinetics and survival. Oncoimmunology 6, e1358331 (2017).

10. Furman, W.L. et al. Improved Outcome in Children With Newly Diagnosed High-Risk Neuroblastoma Treated With Chemoimmunotherapy: Updated Results of a Phase II Study Using hu14.18K322A. J Clin Oncol 40, 335–344 (2022).

11. Ladenstein, R. et al. Investigation of the Role of Dinutuximab Beta-Based Immunotherapy in the SIOPEN High-Risk Neuroblastoma 1 Trial (HR-NBL1). Cancers (Basel) 12 (2020).

12. Anghelescu, D.L. et al. Comparison of pain outcomes between two anti-GD2 antibodies in patients with neuroblastoma. Pediatr Blood Cancer (2014).

13. Navid, F. et al. Phase I trial of a novel anti-GD2 monoclonal antibody, Hu14.18K322A, designed to decrease toxicity in children with refractory or recurrent neuroblastoma. J Clin Oncol 32, 1445–1452 (2014).

14. Dunleavy, K. et al. Dose-adjusted EPOCH-R (etoposide, prednisone, vincristine, cyclophosphamide, doxorubicin, and rituximab) in untreated aggressive diffuse large B-cell lymphoma with MYC rearrangement: a prospective, multicentre, single-arm phase 2 study. Lancet Haematol 5, e609–e617 (2018).

15. Parikh, A.R. et al. Efficacy and Safety of Cetuximab Dosing (biweekly vs weekly) in Patients with KRAS Wild-type Metastatic Colorectal Cancer: A Meta-analysis. Oncologist 27, 371–379 (2022).

16. Mody, R. et al. Irinotecan-temozolomide with temsirolimus or dinutuximab in children with refractory or relapsed neuroblastoma (COG ANBL1221): an open-label, randomised, phase 2 trial. Lancet Oncol 18, 946–957 (2017).

17. Desai, A.V. et al. Outcomes Following GD2-Directed Postconsolidation Therapy for Neuroblastoma After Cessation of Random Assignment on ANBL0032: A Report From the Children’s Oncology Group. J Clin Oncol 40, 4107–4118 (2022).

18. Dong, C. et al. Fcgamma receptor IIIa single-nucleotide polymorphisms and haplotypes affect human IgG binding and are associated with lupus nephritis in African Americans. Arthritis Rheumatol 66, 1291–1299 (2014).

19. Castriconi, R. et al. Identification of 4Ig-B7-H3 as a neuroblastoma-associated molecule that exerts a protective role from an NK cell-mediated lysis. Proc Natl Acad Sci U S A 101, 12640–12645 (2004).

20. Sorkin, L.S. et al. Anti-GD(2) with an FC point mutation reduces complement fixation and decreases antibody-induced allodynia. Pain 149, 135–142 (2010).

21. Ahmed, M. & Cheung, N.K. Engineering anti-GD2 monoclonal antibodies for cancer immunotherapy. FEBS Lett 588, 288–297 (2014).

22. Ladenstein, R. et al. Ch14.18 antibody produced in CHO cells in relapsed or refractory Stage 4 neuroblastoma patients: a SIOPEN Phase 1 study. MAbs 5, 801–809 (2013).

23. Cheung, N.K., Guo, H., Hu, J., Tassev, D.V. & Cheung, I.Y. Humanizing murine IgG3 anti-GD2 antibody m3F8 substantially improves antibody-dependent cell-mediated cytotoxicity while retaining targeting in vivo. Oncoimmunology 1, 477–486 (2012).

24. Federico, S.M. et al. A Pilot Trial of Humanized Anti-GD2 Monoclonal Antibody (hu14.18K322A) with Chemotherapy and Natural Killer Cells in Children with Recurrent/Refractory Neuroblastoma. Clin Cancer Res 23, 6441–6449 (2017)

25. Zhu, Y., Shen, R., Hao, R., Wang, S. & Ho, M. Highlights of Antibody Engineering and Therapeutics 2019 in San Diego, USA: Bispecific Antibody Design and Clinical Applications. Antib Ther 3, 146–154 (2020).

26. Lode, H.N. et al. Vaccination with anti-idiotype antibody ganglidiomab mediates a GD(2)-specific anti-neuroblastoma immune response. Cancer Immunol Immunother 62, 999–1010 (2013).

27. Majzner, R.G. et al. CAR T Cells Targeting B7-H3, a Pan-Cancer Antigen, Demonstrate Potent Preclinical Activity Against Pediatric Solid Tumors and Brain Tumors. Clin Cancer Res 25, 2560–2574 (2019).

28. Yu, A.L. et al. Antibody-dependent cellular cytotoxicity (ADCC) in COG ANBL0032: A phase III randomized trial of chimeric anti-GD2 and GM-CSF/IL2 in high risk neuroblastoma following myeloablative therapy and autologous stem cell transplant (ASCT). Journal of Clinical Oncology 22, 2582–2582 (2004).

29. Wang, W., Erbe, A.K., Hank, J.A., Morris, Z.S. & Sondel, P.M. NK Cell-Mediated Antibody-Dependent Cellular Cytotoxicity in Cancer Immunotherapy. Front Immunol 6, 368 (2015).

30. Perez Horta, Z., Goldberg, J.L. & Sondel, P.M. Anti-GD2 mAbs and next-generation mAb-based agents for cancer therapy. Immunotherapy 8, 1097–1117 (2016).

31. Tibbetts, R. et al. Anti-disialoganglioside antibody internalization by neuroblastoma cells as a mechanism of immunotherapy resistance. Cancer Immunology, Immunotherapy 71, 153–164 (2021).

32. Cheung, N.K. et al. FCGR2A polymorphism is correlated with clinical outcome after immunotherapy of neuroblastoma with anti-GD2 antibody and granulocyte macrophage colony-stimulating factor. J Clin Oncol 24, 2885–2890 (2006).

33. Erbe, A.K. et al. FCGR Polymorphisms Influence Response to IL2 in Metastatic Renal Cell Carcinoma. Clin Cancer Res 23, 2159–2168 (2017).

34. Siebert, N. et al. Neuroblastoma patients with high-affinity FCGR2A, -3A and stimulatory KIR 2DS2 treated by long-term infusion of anti-GD2 antibody ch14.18/CHO show higher ADCC levels and improved event-free survival. OncoImmunology 5, e1235108 (2016).

35. Pereira, N.A., Chan, K.F., Lin, P.C. & Song, Z. The “less-is-more” in therapeutic antibodies: Afucosylated anti-cancer antibodies with enhanced antibody-dependent cellular cytotoxicity. MAbs 10, 693–711 (2018).

36. Morris, Z.S. et al. In Situ Tumor Vaccination by Combining Local Radiation and Tumor-Specific Antibody or Immunocytokine Treatments. Cancer Res 76, 3929–3941 (2016).

37. Voeller, J. et al. Combined innate and adaptive immunotherapy overcomes resistance of immunologically cold syngeneic murine neuroblastoma to checkpoint inhibition. J Immunother Cancer 7, 344 (2019).

38. Yuki, N., Yamada, M., Tagawa, Y., Takahashi, H. & Handa, S. Pathogenesis of the neurotoxicity caused by anti-GD2 antibody therapy. J Neurol Sci 149, 127–130 (1997).

39. Alvarez-Rueda, N. et al. A monoclonal antibody to O-acetyl-GD2 ganglioside and not to GD2 shows potent anti-tumor activity without peripheral nervous system cross-reactivity. PLoS One 6, e25220 (2011).

40. Xiao, W.H., Yu, A.L. & Sorkin, L.S. Electrophysiological characteristics of primary afferent fibers after systemic administration of anti-GD2 ganglioside antibody. Pain 69, 145–151 (1997).

41. Deuis, J.R., Dvorakova, L.S. & Vetter, I. Methods Used to Evaluate Pain Behaviors in Rodents. Front Mol Neurosci 10, 284 (2017).

42. Mogil, J.S. Animal models of pain: progress and challenges. Nat Rev Neurosci 10, 283–294 (2009).

43. Marchettini, P., Lacerenza, M., Mauri, E. & Marangoni, C. Painful peripheral neuropathies. Curr Neuropharmacol 4, 175–181 (2006).

44. Deacon, R.M. Burrowing in rodents: a sensitive method for detecting behavioral dysfunction. Nat Protoc 1, 118–121 (2006).

45. Jirkof, P. et al. Burrowing behavior as an indicator of post-laparotomy pain in mice. Front Behav Neurosci 4, 165 (2010).

46. Shepherd, A.J., Cloud, M.E., Cao, Y.Q. & Mohapatra, D.P. Deficits in Burrowing Behaviors Are Associated With Mouse Models of Neuropathic but Not Inflammatory Pain or Migraine. Front Behav Neurosci 12, 124 (2018).

47. Turner, P.V., Pang, D.S. & Lofgren, J.L. A Review of Pain Assessment Methods in Laboratory Rodents. Comp Med 69, 451–467 (2019).

48. Unituxin [package insert]. United Therapeutics. Durham, NC.

49. Featherly, J., Baxter Wojnowicz, S., Steidl, K. & Burgess, J. Lidocaine for dinutuximab-associated pain? A multicenter retrospective observational cohort study. Pediatr Blood Cancer 69, e29653 (2022).

50. Ladenstein, R. et al. Interleukin 2 with anti-GD2 antibody ch14.18/CHO (dinutuximab beta) in patients with high-risk neuroblastoma (HR-NBL1/SIOPEN): a multicentre, randomised, phase 3 trial. Lancet Oncol 19, 1617–1629 (2018).

51. Bishop, M.W. et al. A Phase 1 and pharmacokinetic study evaluating daily or weekly schedules of the humanized anti-GD2 antibody hu14.18K322A in recurrent/refractory solid tumors. MAbs 12, 1773751 (2020).

52. Evers, M. et al. Anti-GD2 IgA kills tumors by neutrophils without antibody-associated pain in the preclinical treatment of high-risk neuroblastoma. J Immunother Cancer 9 (2021).

53. Zirngibl, F. et al. GD2-directed bispecific trifunctional antibody outperforms dinutuximab beta in a murine model for aggressive metastasized neuroblastoma. J Immunother Cancer 9 (2021).

54. Faraj, S. et al. Neuroblastoma chemotherapy can be augmented by immunotargeting O-acetyl-GD2 tumor-associated ganglioside. Oncoimmunology 7, e1373232 (2017).

55. Fleurence, J. et al. Targeting O-Acetyl-GD2 Ganglioside for Cancer Immunotherapy. J Immunol Res 2017, 5604891 (2017).

56. Saleh, M.N. et al. Phase I trial of the murine monoclonal anti-GD2 antibody 14G2a in metastatic melanoma. Cancer Res 52, 4342–4347 (1992).

57. Sorkin, L.S. Antibody activation and immune reactions: potential linkage to pain and neuropathy. Pain Med 1, 296–302 (2000).

58. Mody, R. et al. Irinotecan, Temozolomide, and Dinutuximab With GM-CSF in Children With Refractory or Relapsed Neuroblastoma: A Report From the Children’s Oncology Group. J Clin Oncol 38, 2160–2169 (2020).

59. Furman, W.L. et al. A Phase II Trial of Hu14.18K322A in Combination with Induction Chemotherapy in Children with Newly Diagnosed High-Risk Neuroblastoma. Clin Cancer Res 25, 6320–6328 (2019).

60. Becker, J.C., Varki, N., Gillies, S.D., Furukawa, K. & Reisfeld, R.A. An antibody-interleukin 2 fusion protein overcomes tumor heterogeneity by induction of a cellular immune response. Proceedings of the National Academy of Sciences of the United States of America 93, 7826–7831 (1996).

61. Haraguchi, M. et al. Isolation of GD3 synthase gene by expression cloning of GM3 alpha-2,8-sialyltransferase cDNA using anti-GD2 monoclonal antibody. Proceedings of the National Academy of Sciences 91, 10455 (1994).

62. Silagi, S. Control of Pigment Production in Mouse Melanoma Cells In Vitro - Evocation and Maintenance. Journal of Cell Biology 43, 263-& (1969).

63. Yang, R.K. et al. Intratumoral treatment of smaller mouse neuroblastoma tumors with a recombinant protein consisting of IL-2 linked to the hu14.18 antibody increases intratumoral CD8+ T and NK cells and improves survival. Cancer Immunol Immunother 62, 1303–1313 (2013).

64. Fowler, C.L., Brooks, S.P., Rossman, J.E. & Cooney, D.R. Postoperative immunotherapy of murine C1300-neuroblastoma. J Pediatr Surg 25, 229–237 (1990).

65. Uphoff, C.C. & Drexler, H.G. Detection of Mycoplasma contamination in cell cultures. Curr Protoc Mol Biol 106, 28 24 21–14 (2014).

66. Erbe, A.K., Wang, W., Gallenberger, M., Hank, J.A. & Sondel, P.M. Genotyping Single Nucleotide Polymorphisms and Copy Number Variability of the FCGRs Expressed on NK Cells, vol. 1441. Springer Science: New York, 2016.

67. Erbe, A.K. et al. KIR/KIR-ligand genotypes and clinical outcomes following chemoimmunotherapy in patients with relapsed or refractory neuroblastoma: a report from the Children’s Oncology Group. J Immunother Cancer 11 (2023).

68. Erbe, A.K., Wang, W., Gallenberger, M., Hank, J.A. & Sondel, P.M. Genotyping Single Nucleotide Polymorphisms and Copy Number Variability of the FCGRs Expressed on NK Cells. Methods Mol Biol 1441, 43–56 (2016).

69. Carlson, P.M. et al. Depth of tumor implantation affects response to in situ vaccination in a syngeneic murine melanoma model. J Immunother Cancer 9 (2021).

70. Muralidharan, A. et al. Comparison of Burrowing and Stimuli-Evoked Pain Behaviors as End-Points in Rat Models of Inflammatory Pain and Peripheral Neuropathic Pain. Front Behav Neurosci 10, 88 (2016).

71. Kang, L. et al. CD38-Targeted Theranostics of Lymphoma with (89)Zr/(177)Lu-Labeled Daratumumab. Adv Sci (Weinh) 8, 2001879 (2021).

72. Hidmark, A.S., Nawroth, P.P. & Fleming, T. Analysis of Immune Cells in Single Sciatic Nerves and Dorsal Root Ganglion from a Single Mouse Using Flow Cytometry. J Vis Exp (2017).

