## Supplemental Figures for "Bispecific GD2 x B7-H3 Antibody Improves Tumor Targeting and Reduces Toxicity while Maintaining Efficacy for Neuroblastoma"

### Supplemental Figures and Data

**Supplemental Figure 1.**

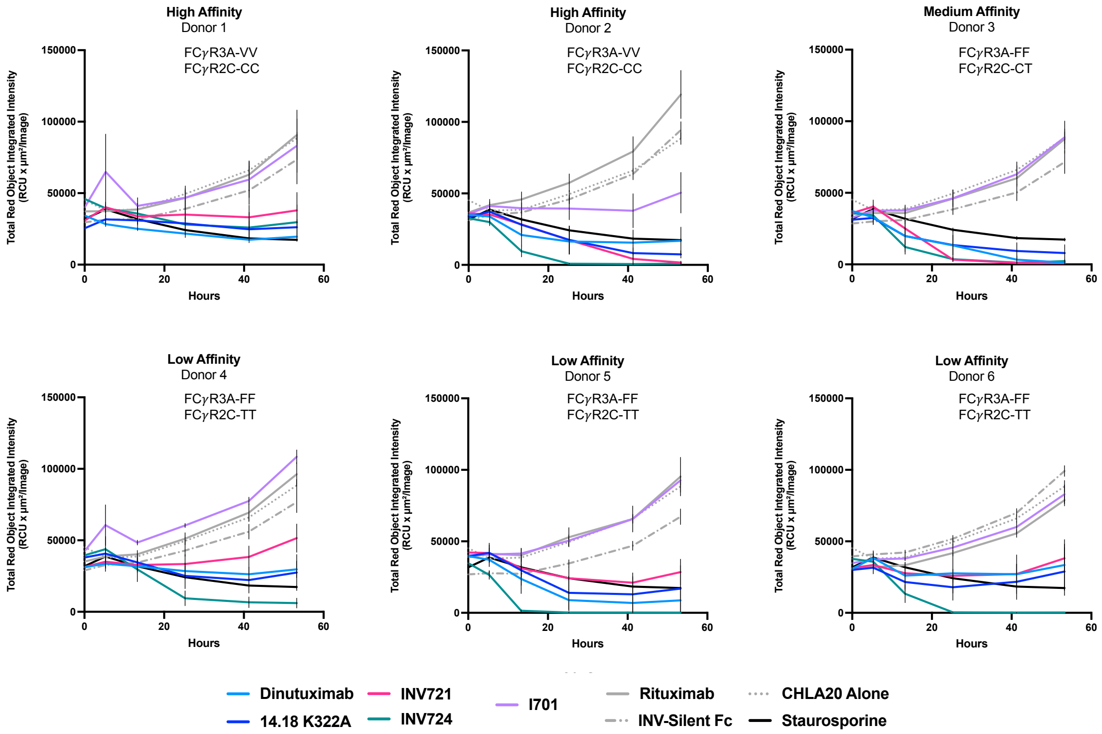

**Supplemental Figure 2.**

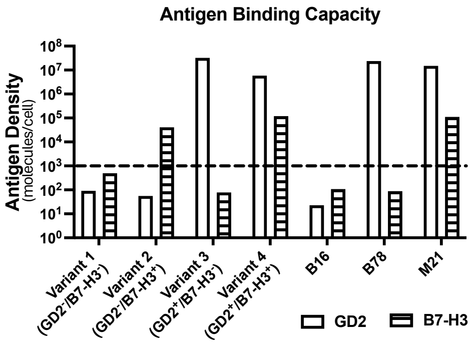

**Supplemental Figure 3.**

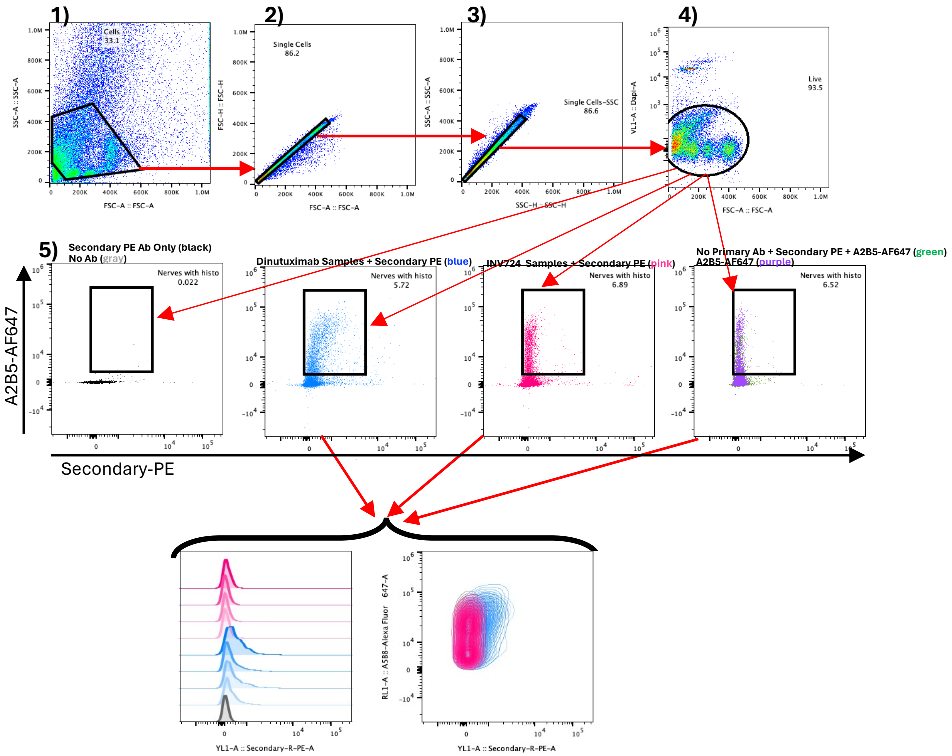


| | Sample Name | Subset Name | Count | Mean : YL1-A |
|---|---|---|---|---|
| | 4 INV724fcs | Nerves with histo | 1127 | 224 |
| | 2 INV724fcs | Nerves with histo | 743 | 179 |
| | 3 INV724.fcs | Nerves with histo | 1410 | 192 |
| | INV724.fcs :s | Nerves with histo | 941 | 183 |
| | 4 Din.fcs | Nerves with histo | 1252 | 550 |
| | 3 Din.fcs | Nerves with histo | 2852 | 410 |
| | 2 Din.fcs | Nerves with histo | 1352 | 522 |
| | 1 Din.fcs | Nerves with histo | 891 | 348 |
| | AF 647 Only.fcs | Nerves with histo | 1017 | 146 |

| | Sample Name | Subset Name | Count | Mean : YL1-A |
|---|---|---|---|---|
| | 4 INV724fcs | Nerves with histo | 1127 | 224 |
| | 2 INV724fcs | Nerves with histo | 743 | 179 |
| | 3 INV724.fcs | Nerves with histo | 1410 | 192 |
| | 1 INV724.fcs | Nerves with histo | 941 | 183 |
| | 4 Din.fcs | Nerves with histo | 1252 | 550 |
| | 3 Din.fcs | Nerves with histo | 2852 | 410 |
| | 2 Din.fcs | Nerves with histo | 1352 | 522 |
| | 1 Din.fcs | Nerves with histo | 891 | 348 |
| | AF 647 Only.fcs | Nerves with histo | 1017 | 146 |

**Supplemental Figure 4.**

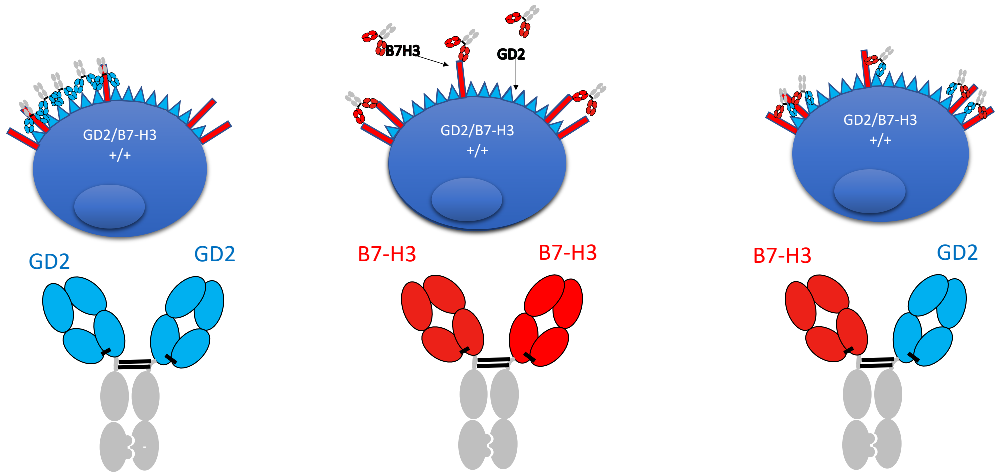
